# R-loops and Topoisomerase 1 facilitate formation of transcriptional DSBs at gene bodies of hypertranscribed cancer genes

**DOI:** 10.1101/2022.12.12.520103

**Authors:** Osama Hidmi, Sara Oster, Jonathan Monin, Rami I. Aqeilan

**Author notes:** **Conflict of Interest:** non.

## Abstract

DNA double-stranded breaks (DSBs) pose a significant threat to genomic integrity, and their generation during essential cellular processes like transcription remains poorly understood. In this study, we employed advanced techniques to map DSBs, R-loops, and Topoisomerase 1 cleavage complex (TOP1cc) and re-analyzed ChIP-seq and DRIP-seq data to comprehensively investigate the interplay between transcription, DSBs, Topoisomerase 1 (TOP1), and R-loops. Our findings revealed the presence of DSBs at highly expressed genes enriched with TOP1 and R-loops, indicating their crucial involvement in transcription-associated genomic instability. Depletion of R-loops and TOP1 specifically reduced DSBs at highly expressed genes, uncovering their pivotal roles in transcriptional DSB formation. By elucidating the intricate interplay between TOP1cc trapping, R-loops, and DSBs, our study provides novel insights into the mechanisms underlying transcription-associated genomic instability. Moreover, we establish a link between transcriptional DSBs and early molecular changes driving cancer development. Notably, our study highlights the distinct etiology and molecular characteristics of driver mutations compared to passenger mutations, shedding light on the potential for targeted therapeutic strategies. Overall, these findings deepen our understanding of the regulatory mechanisms governing DSBs in hypertranscribed genes associated with carcinogenesis, opening avenues for future research and therapeutic interventions.

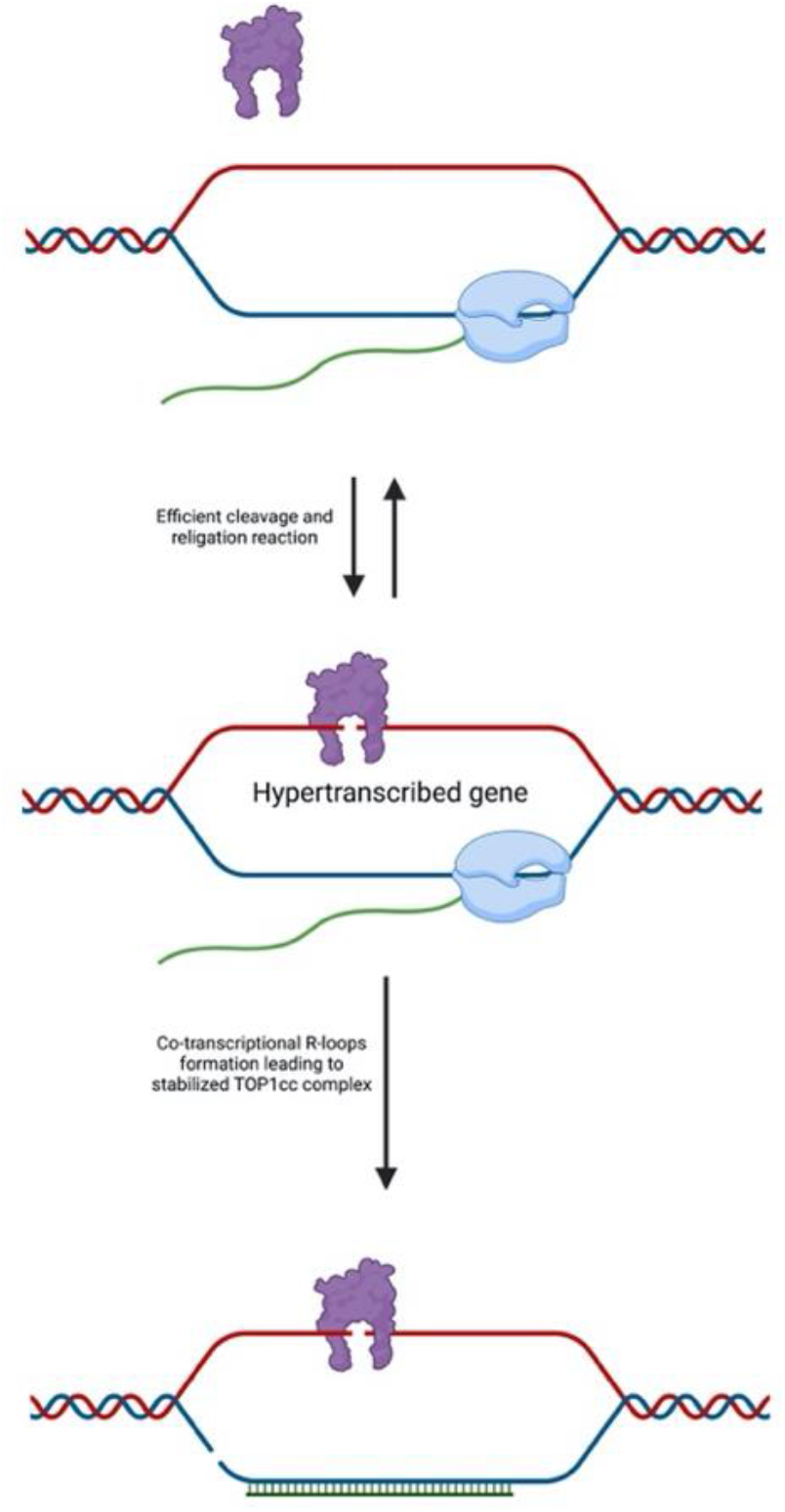

## Introduction

Double-stranded breaks (DSBs) represent the most detrimental form of DNA damage, necessitating prompt repair mechanisms to preserve genome integrity. While cytotoxic levels of DSBs primarily result from external factors such as ionizing radiation and UV radiation, DSBs can also arise naturally during physiological processes like transcription and replication (Wilhelm, Said and Naim, 2020).

In replicating cells, transcription can give rise to DSBs through collisions with the replication machinery, particularly when transcription and replication occur in opposite directions (Mirkin and Mirkin, 2005; Helmrich, Ballarino and Tora, 2011; Hamperl et al., 2017). This phenomenon, commonly referred to as transcription-replication conflict, occurs when DNA polymerases, the main players in replication, and RNA polymerases, the primary components of the transcriptional machinery, encounter each other in close proximity within a gene. Given the heightened transcriptional and replication activity observed in cancer cells, it is reasonable to expect an increased incidence of such conflicts (Helmrich, Ballarino and Tora, 2011; Hamperl et al., 2017). Interestingly, this collision between transcription and replication predominantly occurs at transcription termination sites (TTS) and can be mitigated by the activity of topoisomerase 1 (TOP1) (Promonet et al., 2020). Additionally, elevated transcription levels have been strongly correlated with the occurrence of DSBs (Lensing et al., 2016). Nonetheless, the precise mechanisms by which transcription itself leads to the generation of DSBs remain subjects of ongoing investigation.

During transcription elongation, the DNA strand unwinds around the transcribing RNA Pol II, significantly increasing the likelihood of newly transcribed RNA molecules annealing to their DNA template. This process gives rise to a unique nucleotide structure known as an R-loop, composed of a DNA:RNA hybrid and a displaced single-stranded DNA (Petermann, Lan and Zou, 2022). Short R-loops (<10 bp) play essential roles in various cellular processes and are transiently formed during transcription, promptly resolved by the enzyme RNase H1 (Cerritelli and Crouch, 2009) and RNA-DNA helicases. However, longer R-loops have been implicated in jeopardizing genome integrity by promoting the generation of DSBs. The precise mechanisms underlying the R-loop-mediated DSB formation are still poorly understood. One proposed mechanism suggests that R-loops expose the displaced single-stranded DNA to endonucleases (Sollier and Cimprich, 2015), rendering it more susceptible to single-stranded breaks (SSBs). Furthermore, specific endonucleases such as XPG, XPF, and FEN1 have been found to recognize R-loop structures, leading to cleavage of the single-stranded DNA hybridized with RNA within the R-loop, resulting in SSBs. However, the process by which these SSBs are converted into DSBs remains unknown.

During transcription elongation, the movement of RNA Pol II induces the generation of negative and positive supercoiling behind and ahead of the transcription bubble, respectively (Liu and Wang, 1987; Ma and Wang, 2016). The resolution of these supercoiling events is crucial for maintaining efficient transcription, particularly at highly transcribed genes (Ma and Wang, 2016). Topoisomerase 1 (TOP1) plays a central role in the resolution of both positive and negative supercoiling. Through its catalytic domain, TOP1 introduces transient single-strand breaks in the DNA, allowing for the relaxation of supercoils. Subsequently, the broken DNA strands are re-ligated (Pommier et al., 2016, 2022). Normally, the formation of a transient Topoisomerase cleavage complex (TOP1cc) ensures that the DNA is cut and rapidly rejoined, resulting in no persistent DNA damage. However, if TOP1 becomes trapped in its catalytic state, whether due to physiological or pathological factors, the single-stranded DNA remains nicked, and the TOP1cc complex bound to the DNA needs to be removed through the Tyrosyl-DNA Phosphodiesterase 1 (TDP1) excision pathway for proper repair of the break to occur (Cristini et al., 2016; Pommier et al., 2022).

Mutations are recognized as the primary driving force behind tumorigenesis, leading to the dysregulation of cellular processes and the development of cancer (Stratton, Campbell and Futreal, 2009). The prevailing consensus is that these mutations arise spontaneously during physiological conditions, and the observed bias in the mutational profile of cancer cells obtained from patients is attributed to the selective advantage conferred by specific mutations through the process of Darwinian evolution(Greaves and Maley, 2012; McGranahan and Swanton, 2017). Functionally significant mutations, known as “driver mutations,” occur in genes referred to as “driver genes,” whereas non-functional frequent mutations are classified as “passenger mutations”. Computational tools have been developed to distinguish driver mutations from passenger mutations based on their deviation from random background mutations and their downstream effects (Nik-Zainal et al., 2016; Martínez-Jiménez et al., 2020; Malebary and Khan, 2021). Despite these advancements, our understanding of the underlying etiology of cancer mutations, particularly the differences between mutations that contribute to cancer fitness and those that do not, remains limited.

To detect DNA DSBs, various methods have been employed. Antibodies against DNA damage markers such as anti-γ-H2AX have proven valuable in assessing changes in DNA damage under different treatments and conditions. However, while these antibodies provide valuable information regarding DNA damage, they do not offer precise localization of the break sites within the genome. Chromatin immunoprecipitation sequencing (ChIP-seq) assays utilizing such antibodies can help identify the genomic regions associated with these breaks. Nonetheless, due to the broad distribution of γ- H2AX, which can extend over several kilobase pairs around the break site, it may not provide the desired high-resolution detection of the exact break site. In this study, we employed a recently developed technique known as in-suspension break labeling *in situ* and sequencing (sBLISS) (Bouwman et al., 2020) to address these limitations. sBLISS enables the detection of DSBs with single-nucleotide resolution, providing highly precise information about the location of breaks across the genome. Notably, sBLISS exhibits excellent discrimination between DSBs occurring at proximal genomic locations, thereby enhancing our ability to study transcriptional DSBs. Moreover, the effectiveness of sBLISS in detecting physiological DSBs has been confirmed previously (Hazan et al., 2019), further highlighting its potency as a powerful tool for investigating transcriptional DSBs.

In this study, we employed a comprehensive approach utilizing multiple omic techniques, namely sBLISS, ChIP-seq, and DRIP-seq, to elucidate the intricate mechanism underlying the generation of physiological DSBs during transcription. Through the integration of these complementary methodologies, we uncovered a novel mechanism involving the interplay of transcription, R-loops, TOP1, and TOP1cc in facilitating the formation of DSBs at active genomic regions, particularly within highly transcribed oncogenes.

## Results

### Physiological breaks are enriched at highly transcribed genes

In our previous work, we observed enrichment of physiological DSBs in active genomic regions such as promoters, insulators, and super-enhancers using BLISS (Hazan et al., 2019). To further unravel the underlying mechanism governing physiological DSBs, we employed the recently developed sBLISS method (Bouwman et al., 2020) to comprehensively map the distribution of DSBs across the genome in untreated MCF-7 breast cancer luminal cells. Comparative analysis of observed breaks with expected breaks in various chromatin states revealed a significantly higher observed-to-expected ratio at active promoters, strong enhancers, transcription elongation sites, and transcription transition regions, while repressed regions exhibited a considerably lower ratio (Figure 1A). This heightened enrichment of breaks in active regions, compared to the expected distribution, further substantiates the association between transcription and the generation of DSBs in these genomic loci. To further validate the relationship between transcription and DSBs, we quantified gene expression using the CEL-seq RNA sequencing technique (Hashimshony et al., 2012), and correlated it with the corresponding break density for each gene. Interestingly, our analysis, incorporating both gene expression data we generated and previously published RNA-seq data for MCF-7 cells, unveiled a positive correlation between gene expression and break density (Figure 1B and C, Supplementary Figure 1A, B, and D-K), suggesting that highly transcribed genes exhibit a propensity for increased DSB density. Additionally, gene ontology analysis of the top 400 genes with the highest number of breaks in MCF-7 cells revealed enrichment in biological processes such as translation, gland morphogenesis, and response to estradiol, which are known to be highly active in breast epithelial tissue (Supplementary Figure 1C). This finding further underscores the association between active genes and elevated break densities.

**Figure 1:**
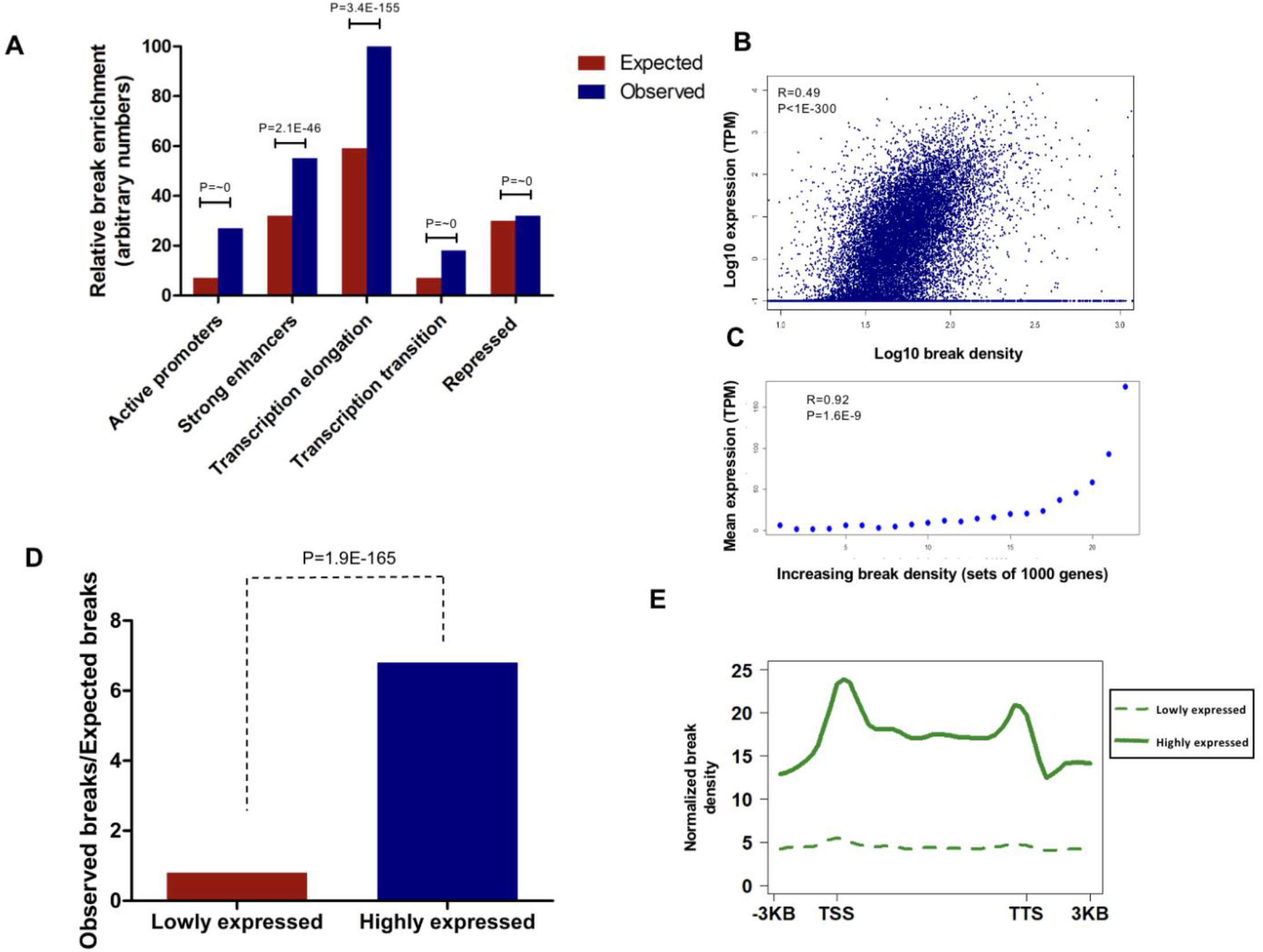
Breaks are enriched at highly expressed genes. **A**. The distribution of DSBs in breast cancer MCF7 cells along ChromHMM-defined chromatin states of HMEC. Bar height represents break enrichment relative to other chromatin states. **B**. A zoomed log scaled scatter plot showing a positive correlation between break density and expression levels; each dot represents a gene, and for each gene break density and expression were measured and normalized to gene size. **C**. Mean expression (TPM) positively correlates with break density, genes were grouped into 22 groups with increasing break enrichment, the 22nd group being the group with the highest mean break enrichment. **D**. Ratio of observed breaks vs expected breaks at highly and low expressed genes. **E.** Plot showing break density at gene body of the 1000 most transcribed genes (continuous line) and 1000 least transcribed genes (dashed line). sBLISS experiments were done in duplicates, and CEL-seq was done in quadruplicates.

To investigate the nature of these transcriptional DSBs, we selected the top 1000 highly expressed genes and the bottom 1000 lowly expressed genes based on CEL-seq data and calculated the ratio of observed to expected breaks for each gene. Consistent with our predictions, highly expressed genes exhibited a substantially higher ratio (∼7-fold) compared to lowly expressed genes (<1) (Figure 1D). Moreover, when analyzing the distribution of break densities across the gene body of these genes (Figure 1E), we observed an elevated break density at the gene body of highly transcribed genes in comparison to lowly transcribed genes, with prominent peaks at the transcription start site (TSS) and transcription termination site (TTS).

To address the possibility that the observed DSBs are a consequence of replication and/or transcription-replication collisions, we replicated the experiment using MCF-7 cells synchronized in the G1 phase of the cell cycle (Supplementary Figure 2A, B). Remarkably, we obtained similar results (Supplementary Figure 2C and D), further substantiating the transcription-dependent nature of these DSBs. These data clearly suggest that physiological DSBs are more enriched at active regions, specifically at highly transcribed genes, and point towards transcription as a direct or indirect cause of physiological DSBs.

### Transcriptional DSBs are associated with TOP1

The presence of TOP1 at regulatory elements such as insulators, promoters, and enhancers, particularly those exhibiting high break density, has been previously reported (Baranello et al., 2016; Hazan et al., 2019). To directly assess the involvement of TOP1 in transcriptional DSB formation, we examined the correlation between TOP1 levels and gene expression using our previously published TOP1 ChIP-seq data (Hazan et al., 2019). Remarkably, we observed a positive correlation between TOP1 levels and the expression of moderately and highly expressed genes (Figure 2A, B, Supplementary Figure 3A). Additionally, we found a significant enrichment of breaks at TOP1 peaks throughout the genome (Figure 2C). Furthermore, when comparing the ratio of observed to expected breaks, high TOP1-enriched genes exhibited a substantially higher ratio (∼12-fold) compared to low TOP1-enriched genes (Figure 2D). To further validate the significance of TOP1, we performed TOP1 knockdown using siRNA (Supplementary Figure 3B) and evaluated the impact on DNA breaks using immunofluorescence staining. As depicted in Figure 2E and F, TOP1-depleted MCF7 cells displayed a significant decrease in γ-H2AX and 53BP1 foci per nucleus compared to siRNA control transfected cells. These findings collectively underscore the crucial role of TOP1 in transcriptional DSB formation.

**Figure 2:**
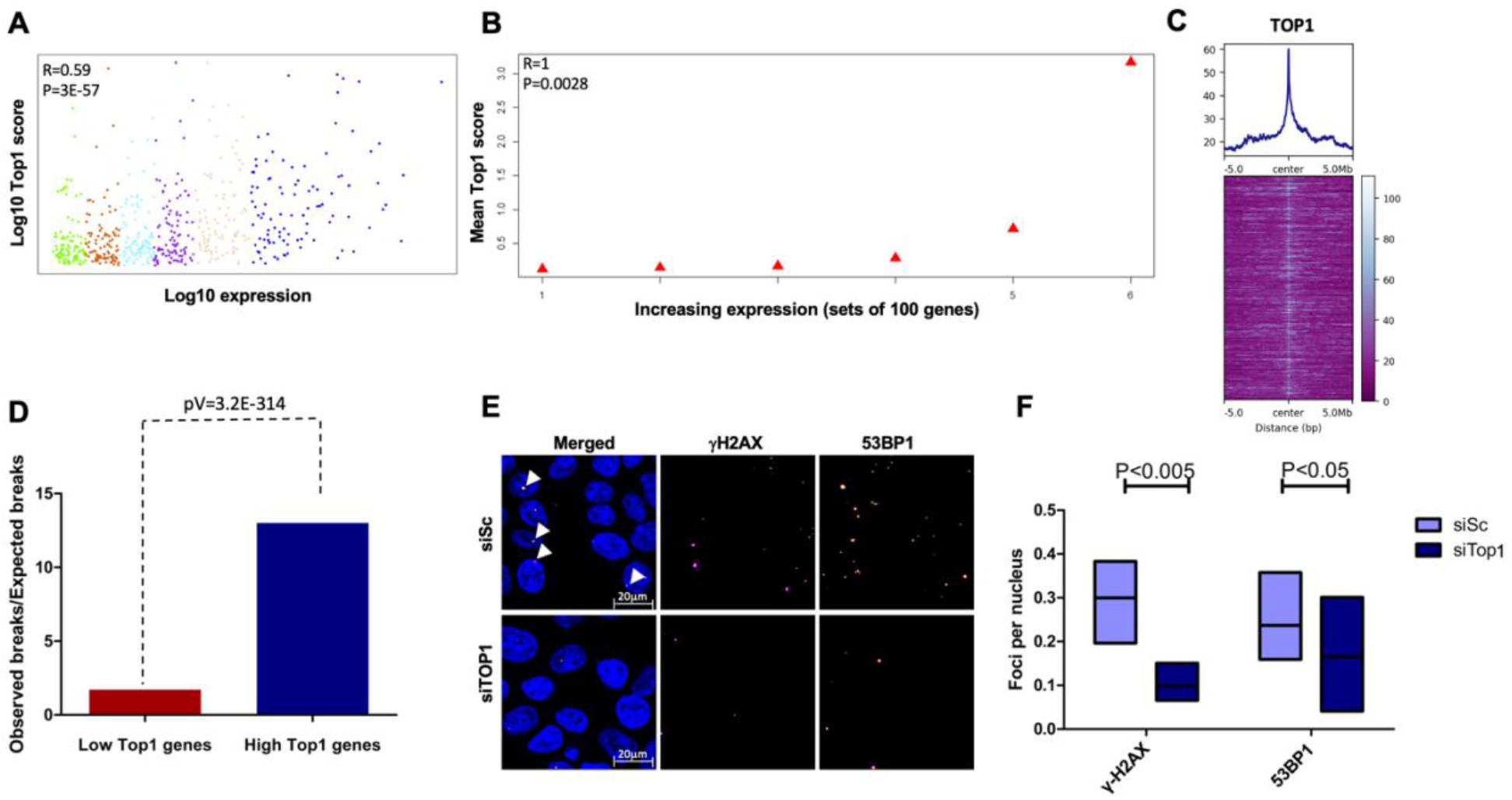
Transcriptional DSBs are associated with TOP1. **A.** A log scaled scatter plot showing positive correlation between expression and TOP1 levels for the top 50% expressed genes; each dot represents a gene, and each color represents a gene group used in **B**. **B.** Median of expression (TPM) correlates with TOP1 levels, for the top 50% expressed genes. **C.** Heatmap of break density at TOP1 binding regions across the genome, the plot shows DSBs at the center of TOP1 peaks. **D.** Ratio of observed breaks to expected breaks for low TOP1 and high TOP1 genes. **E.** Representative images of immunofluorescent staining of MCF-7 ctrl (siSc) and TOP1-depleted (siTOP1) MCF-7 cells using γ-H2AX and 53BP1 antibodies. **F.** Quantification of the number of γ-H2AX and 53BP1 foci per nucleus in MCF-7 ctrl and MCF- 7 siTOP1 (E). sBLISS and immunofluorescence experiments were done in duplicates.

### Transcriptional DSBs are associated with R-loops

R-loops, known for their detrimental impact on genome integrity, are facilitated by the transcriptional machinery. Given the link between TOP1 and R-loops, which was further supported by correlating TOP1 levels with R-loop levels (Supplementary Figure 3C), we proceeded to investigate the role of R-loops in transcriptional DSB formation. Correlating R-loop levels, using published DRIP-seq data from untreated MCF-7 cells (GSE81851), with gene expression revealed a positive correlation in the top 50% expressed genes (Figure 3A and B, Supplementary Figure 3D). Mapping break enrichment at R-loop peaks demonstrated an elevated break enrichment at these sites (Figure 3C). Furthermore, comparing the observed to expected ratio of breaks between genes enriched or depleted in R-loops showed that R-loop-enriched genes had a higher ratio (∼9-fold) compared to those lacking R-loops (Figure 3D).

**Figure 3:**
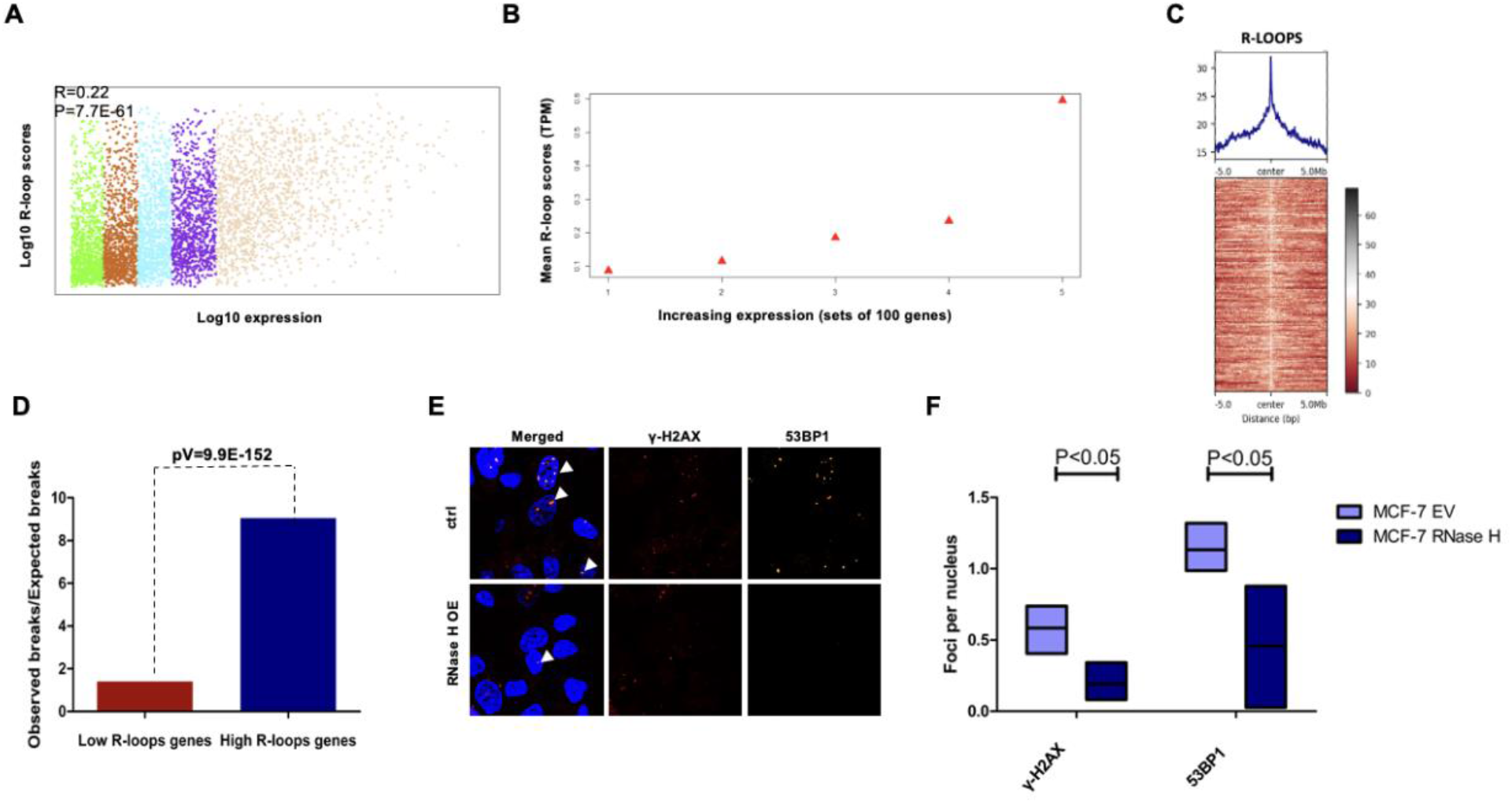
Transcriptional DSBs are associated with R-loops. **A**. A log scaled scatter plot showing positive correlation between expression and R-loops levels for the top 50% expressed genes, each dot represents a gene. DRIP data was extracted from GSE81851. **B.** Mean expression (TPM) correlates with R-loops levels for the top 50% expressed genes. **C.** Heatmap of break density at R-loops regions across the genome, the plot shows DSBs at the center of TOP1 peaks. **D.** Ratio of observed breaks to expected breaks for low R-loops and high R-loops genes. **E.** Representative images of immunofluorescent staining of MCF-7 EV and MCF-7 overexpressing RNase H1 for γ-H2AX and 53BP1. **F.** Number of γ-H2AX and 53BP1 foci per nucleus, for MCF-7 EV and MCF-7 RNase H1. sBLISS and immunofluorescence experiments were done in duplicates.

To investigate the impact of R-loop depletion on global DSBs, we overexpressed RNase H1, an enzyme responsible for degrading the RNA component of R-loops (Cerritelli and Crouch, 2009), in MCF-7 cells and performed immunofluorescence staining for 53BP1 and γ-H2AX (Figure 3E, F). Intriguingly, cells overexpressing RNase H1 displayed fewer γ-H2AX and 53BP1 foci per nucleus compared to control cells, indicating the involvement of R-loops in the formation of physiological DSBs.

### TOP1 knockdown and RNase H1 overexpression decrease break enrichment specifically at the gene body of highly transcribed genes

To unravel the intricate involvement of TOP1 and R-loops in the formation of transcriptional DSBs, we conducted two different manipulations in MCF-7 cells. Our investigations centered around depleting TOP1 in cells overexpressing RNase H1, as well as control cells (Supplementary Figure 4A-C). By utilizing the sBLISS technique, we meticulously mapped the changes in DSBs across various genomic regions.

In our initial analysis, we explored break enrichment in distinct chromatin states and made interesting observation. Specifically, regions associated with transcription elongation and transcription transition chromatin states exhibited a notable reduction in break frequency in the manipulated samples compared to the control group (Figure 4A). This finding prompted us to further investigate the impact of RNase H1 overexpression and TOP1 knockdown on highly active regions, with a particular focus on the top 1000 expressed genes identified through CEL-seq analysis of these MCF-7 cells.

**Figure 4:**
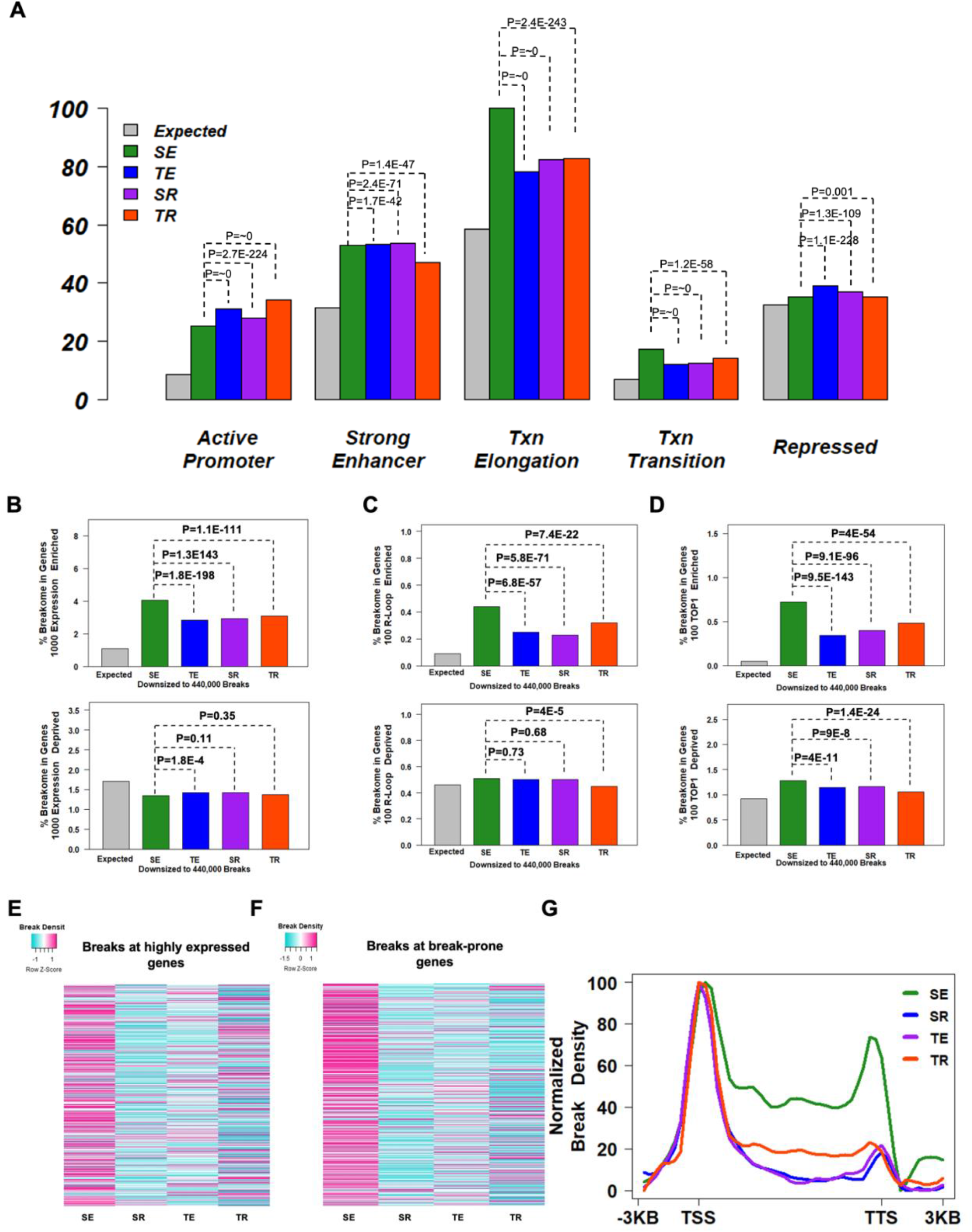
RNase H1 overexpression and TOP1 knockdown decrease break enrichment at gene body of highly expressed genes. **A.** The distribution of DSBs in breast cancer MCF7 cells along ChromHMM- defined chromatin states of HMEC in control cells (SE), cells knocked down for TOP1 (TE), cells overexpressing RNase H1 (SR), and cells knocked down for TOP1 and overexpressing RNase H1 (TR). Bar height is break enrichment relative to other chromatin states. **B**. Breakome percentage in 1000 most transcribed genes (top) and 1000 least transcribed genes (bottom) with the different manipulations. Expression data are taken from our CEL-seq. **C**. Breakome percentage in 100 top R-loop enriched genes (top) and 100 top R-loop deprived genes (bottom) with the different manipulations. **D**. Breakome percentage in 100 top TOP1 enriched genes (top) and 100 top TOP1 deprived genes (bottom) with the different manipulations. **E.** color-coded heat map representing the change in break density for the top 1000 expressed genes. The break density values have been standardized using z-scores for rows to enhance the visualization of relative differences. **F.** color-coded heat map representing the change in break density for the top 2000 break prone genes. **G**. Plot showing break density at gene body of highly transcribed genes shown in **B** with the different manipulations. sBLISS experiments were done in duplicates.

Remarkably, our investigation revealed a consistent pattern in highly transcribed genes. These genes displayed a marked decrease in DSB enrichment upon TOP1 depletion and/or RNase H1 overexpression (Figure 4B, E and Supplementary Figure 4D). In contrast, we observed no significant effect on the break density of low-expressed genes (Figure 4B, Supplementary Figure 4D). Furthermore, genes enriched with R-loops exhibited a substantial reduction in break enrichment following RNase H1 overexpression and/or TOP1 knockdown, while genes lacking R-loops showed no significant change (Figure 4C). Similarly, genes enriched with TOP1 exhibited a significant decrease in break density compared to genes devoid of TOP1 (Figure 4D).

Of particular interest, the impact of TOP1 knockdown and/or RNase H1 overexpression was particularly pronounced in break-prone genes. We defined break-prone genes as the top 2000 genes exhibiting the highest break enrichment in the untreated sample, and intriguingly, both TOP1 knockdown and RNase H1 overexpression led to a significant decrease in break enrichment in these genes (Figure 4F).

We delved deeper into the distribution of breaks along the gene body upon TOP1 depletion and/or RNase H1 overexpression. Strikingly, our analysis demonstrated a consistent reduction in break density within the gene body and transcription termination site (TTS) of highly expressed genes, while the break density at the transcription start site (TSS) remained largely unaffected (Figure 4G and Supplementary Figure 4E). These findings provide strong evidence supporting the notion that the vulnerability of gene bodies in highly transcribed genes is mediated by R-loops and TOP1. Moreover, our observations align with previous reports suggesting the catalytic activity of TOP1 specifically at gene bodies, rather than the TSS (Baranello et al., 2016).

### Fragility of estradiol responsive genes is mediated by R-loops and TOP1

To investigate the impact of increased transcription on global DNA damage, MCF-7 cells were treated with estradiol (E2) to induce transcription. Immunofluorescence staining for γ-H2AX and 53BP1 was performed to assess DNA damage levels (Figure 5A, B). As expected, cells treated with E2, exhibited a higher number of 53BP1 and γ-H2AX foci per nucleus. However, this increase in DNA damage was suppressed when RNase H1 was overexpressed or TOP1 was depleted, indicating that R-loops and TOP1 mediate the increase in DNA damage upon E2 treatment.

**Figure 5:**
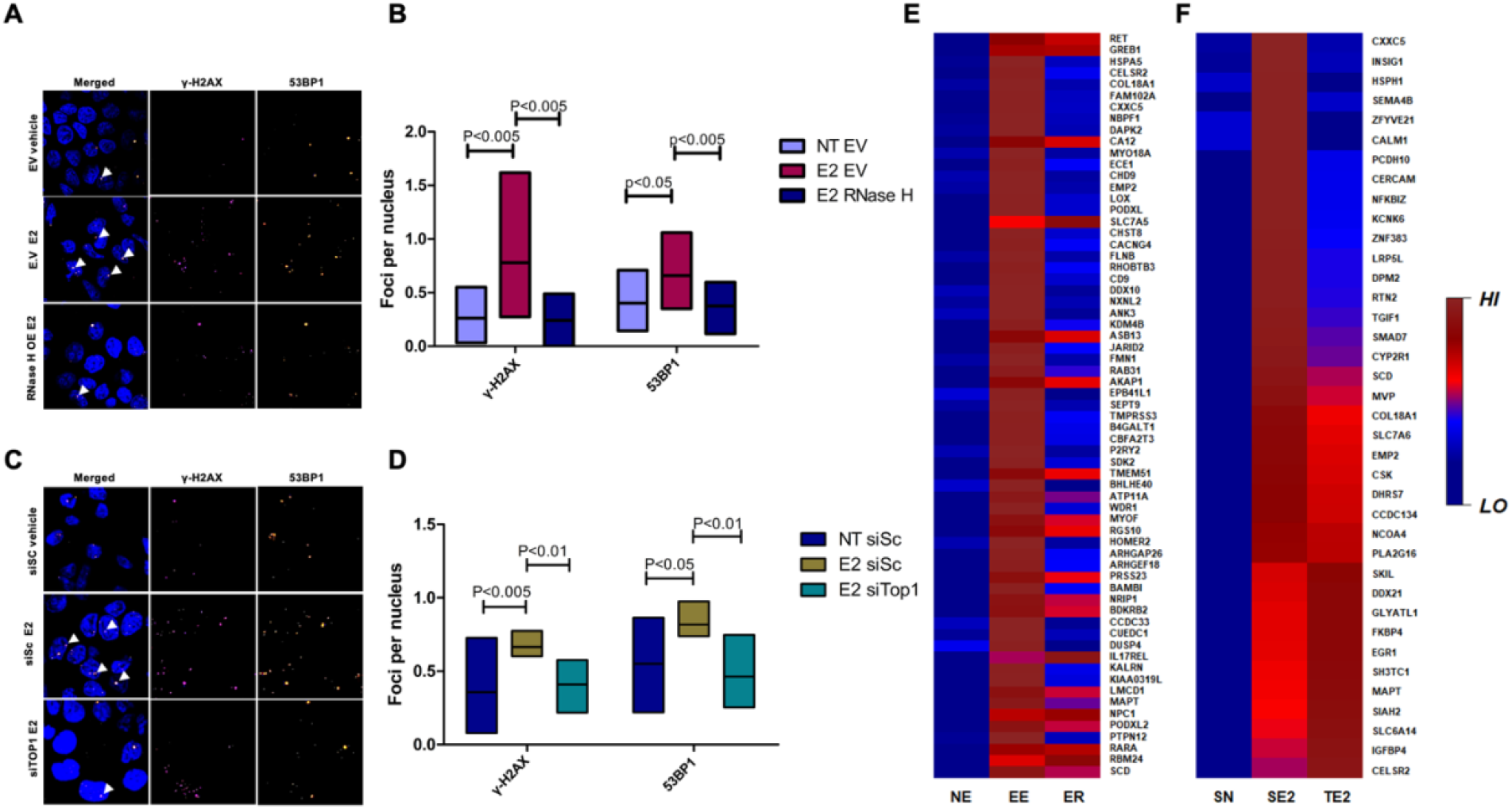
Estradiol associated DSBs are mediated by R-loops and TOP1. **A.** Representative images of immunofluorescent staining of MCF-7 EV non treated, MCF-7 EV treated with estradiol (E2), and MCF-7 cells overexpressing RNase H1 and treated with E2, for γ-H2AX and 53BP1. **B.** Quantification of number of γ-H2AX and p53BP1 foci per nucleus, for MCF-7 EV NT and MCF-7 EV treated with E2, and MCF-7 cells overexpressing RNase H1 treated with E2. **C.** Representative images of immunofluorescent staining of MCF-7 siSc non treated, MCF-7 siSc treated with estradiol (E2), and MCF-7 cells knocked down for TOP1 and treated with E2, for γ-H2AX and 53BP1. **D.** Quantification of number of γ-H2AX and p53BP1 foci per nucleus, for MCF-7 siSc NT and MCF-7 siSc treated with E2, and MCF-7 cells knocked down for TOP1 and treated with E2. **E.** Color coded heatmap showing the change in break density between control (NE), cells incubated with estradiol (EE), cells incubated with estradiol and overexpressing RNase H1 (ER), for estrogen responsive genes with the highest positive differential break density. **F.** Color coded heatmap showing the change in break density between control (SN), cells incubated with estradiol (SE2), cells knocked down for TOP1 (TE2), for estrogen responsive genes with the highest positive differential break density. sBLISS and immunofluorescence experiments were done in duplicates.

To further investigate these findings, sBLISS was used to map DSBs in different conditions: control cells (NE/SN), cells treated with E2 (EE/SE2), and cells overexpressing RNase H1 or knocked down for TOP1 and treated with E2 (ER/TE2).

Most of estrogen-responsive genes, which exhibited a significant increase in expression upon E2 treatment (extracted from GSE27463) (Hah et al., 2011), displayed higher break density upon E2 treatment as opposed to down-regulated genes (Supplementary Figure 5 A-D). Importantly, most of these genes showed decreased break density when R-loops or TOP1 were depleted (ER/TE2) compared to estradiol treatment alone, indicating that the increase in transcriptional DSBs upon E2 treatment is mediated by R-loops and TOP1.

Moreover, among the estrogen-responsive genes that exhibited increased break density following estradiol induction, the fragility of these genes was significantly reduced by depleting R-loops or TOP1 (Figure 5 E and F), further confirming the involvement of TOP1 and R-loops in transcriptional DSBs, particularly in the context of estrogen-associated DSBs. Additionally, pathway analysis of these estrogen-responsive genes using DisGenNET (Piñero et al., 2021) revealed enrichment in tumor progression, neoplasm metastasis, breast carcinoma, and malignant neoplasm of the breast (Supplementary Figure 5E), suggesting that estrogen treatment upregulates breast cancer-associated genes, leading to DSB formation at these genes through R-loops and TOP1.

Taken together, these findings demonstrate that increased transcription results in elevated DSB formation, and this transcriptional DSBs are associated with the presence of R-loops and TOP1.

### DSBs at highly transcribed genes might limit transcription

To ensure the validity of the observed decrease in break enrichment at highly expressed genes, we meticulously investigated any potential effects on transcription levels caused by the manipulations. Employing CEL-seq, we analyzed the change in normalized expression counts of the top 1000 highly expressed genes under different manipulations (Supplementary Figure 6A, B). Remarkably, cells overexpressing RNase H1 (SR) or depleted of TOP1 (TE) displayed a significant increase in expression levels for the top 1000 expressed genes, while the combined manipulation (TR) did not exhibit the same effect (Supplementary Figure 6C-G). Furthermore, a comparative analysis of gene sets sorted by expression in the control sample revealed intriguing insights. Specifically, the highest expressed group (group 1) demonstrated the largest percentage of genes with increased transcription upon manipulations (Supplementary Figure 6H). This observation was further corroborated by examining the normalized expression counts of these gene sets, where the highest expressed group exhibited the most substantial increase in expression levels (Supplementary Figure 6I).

Notably, when specifically scrutinizing the change in expression at highly expressed genes that exhibited a decrease in break density, we observed a predominant trend towards increased expression (Supplementary Figure 6J-O). This trend aligns with our understanding that intragenic DSBs are known to impede transcription. However, it is important to note that not all genes demonstrated increased expression in response to decreased DSBs, and the patterns appeared to be variable. These intriguing findings suggest that the effects of TOP1 depletion and/or RNase H1 overexpression on gene expression are context-dependent, exerting diverse influences on different genes. Taken together, our results strongly imply that transcriptional DSBs, facilitated by the interplay between TOP1 and R-loops, may act as impediments to transcription at highly expressed genes, shedding light on the intricate relationship between DNA breakage, transcriptional regulation, and genomic stability.

### Interplay between TOP1cc and R-loops in Transcriptional DSBs Formation

To gain deeper insights into the mechanisms underlying transcriptional DSBs formation, we investigated the interplay between TOP1cc and R-loops at highly expressed genes. Specifically, we aimed to determine whether TOP1cc promotes R-loop formation or, conversely, whether R-loops enhance the trapping events of TOP1cc.

To address these questions, we performed DNA-RNA immunoprecipitation sequencing (DRIP-seq) experiments following TOP1 knockdown and/or overexpression of RNase H1. Consistent with the previous analysis of published DRIP-seq, our analysis revealed a positive correlation between R-loop levels and gene expression, as illustrated in Supplementary Figure 7A. Examining the changes in R-loop levels resulting from the different manipulations, we observed a significant increase in R-loops at highly expressed genes following TOP1 KD (Supplementary Figure 7B-D). These findings align with previous reports demonstrating the role of TOP1 in preventing R-loop accumulation (Li et al., 2015; Manzo et al., 2018). Conversely, examining the effect of RNase H1 OE at highly expressed genes showed a decrease in DRIP signal, indicating a reduction in R-loops (Supplementary Figure 7E-G). Remarkably, when selecting genes exhibiting decreased breaks upon RNase H1 OE, we observed a concomitant decrease in R-loop levels (Supplementary Figure 7H). These results further confirm the role of TOP1 in preventing R-loop accumulation, although we do not exclude the possibility that trapping of TOP1cc may contribute to increased R-loop formation, as previous studies have shown that treatment with the trapping agent CPT leads to increased R-loops at gene bodies (Cristini et al., 2019).

To investigate the potential scenario of TOP1cc trapping by R-loops, we performed chromatin immunoprecipitation sequencing (ChIP-seq) using an antibody specifically recognizing the bond between TOP1 and DNA in TOP1cc, following TOP1 KD and/or RNase H1 OE. The analysis revealed a positive correlation between TOP1cc and TOP1 at the top 50% expressed genes (Figure 6A). Additionally, genes enriched with TOP1cc exhibited higher levels of TOP1 (Figure 6B), suggesting that TOP1 binding to DNA is associated with the formation of TOP1cc. This association implies that the detected TOP1 binding sites are likely actively involved in DNA cleavage and re-ligation reactions, ultimately resulting in TOP1cc formation. Furthermore, genes enriched with TOP1cc showed a significantly higher enrichment of DSBs compared to genes depleted of TOP1cc (Figure 6C), underscoring the detrimental effect of TOP1cc on genomic integrity.

**Figure 6.**
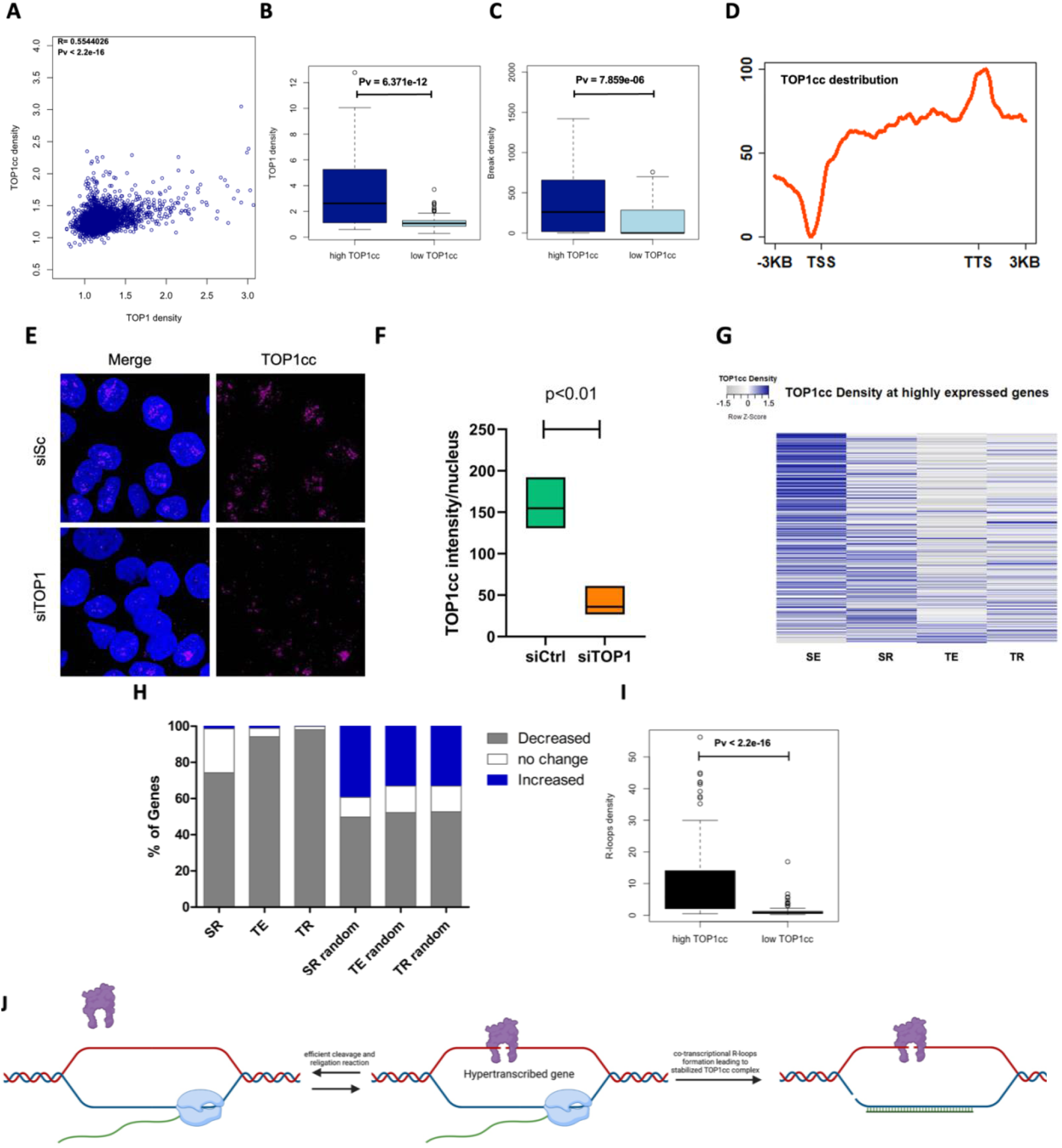
TOP1 KD and/or RNase H OE decrease TOP1cc at highly expressed genes. **A.** Scatter plot showing the correlation between TOP1 and TOP1cc density at the top 50% expressed genes. **B.** Boxplot showing TOP1 density mean at high TOP1cc genes vs low TOP1cc genes. **C.** Boxplot showing break density at high TOP1cc genes vs low TOP1cc genes. **D.** distribution of TOP1cc along the gene bodies of highly expressed genes. **E.** Representative images of immunofluorescence staining of TOP1cc after TOP1 KD. **F.** Quantification of TOP1cc intensity. **G.** Color coded heatmap showing the change in TOP1cc density with the different manipulations at highly expressed genes. H. stacked bar plot showing the percentage of genes the exhibited a decrease/no change/ increase in TOP1cc density for highly expressed genes that decreased in break density upon the different manipulations. **I.** Boxplot showing R-loops density mean at high TOP1cc genes vs low TOP1cc genes. **J.** Graphical illustration of the proposed mechanism of transcriptional DSBs, optimally, TOP1 participates in efficient cleavage and re-ligation reactions maintaining the native structure of DNA and preventing genomic instability. When co-transcriptional R-loops are formed, the engaged TOP1 have higher chance of trapping in its TOP1cc form, leaving a single stranded break (SSB) behind. Along with the SSB formed as a result of R-loops processing, a DSB is formed during transcription.

The distribution of TOP1cc along the gene bodies of highly expressed genes were consistent with previous mapping of TOP1 catalytic activity (Figure 6D), further validating our approach, and showing that TOP1cc can be trapped at physiological levels during hypertranscription.

TOP1 depletion resulted in decreased global physiological TOP1cc levels as observed in immunofluorescence staining (Figure 6E and F). Examining the changes in TOP1cc levels at highly expressed genes upon TOP1 KD revealed a depletion of TOP1cc (Figure 6G, Supplementary Figure 7I and J). Notably, RNase H1 OE also led to a decrease in TOP1cc at highly expressed genes, suggesting a potential role for R-loops in TOP1cc trapping. To further investigate this, we focused on genes exhibiting decreased DSBs following each manipulation. Strikingly, RNase H1 OE resulted in a decrease in TOP1cc at the majority of highly expressed genes displaying decreased DSBs, with a more pronounced effect observed upon TOP1 depletion with both TOP1 KD and RNase H1 OE yielding the highest proportion of genes with decreased TOP1cc (Figure 6H and Supplementary Figure 7K-P). Collectively, these results highlight the involvement of TOP1cc trapping at physiological conditions in transcriptional DSB formation and demonstrate that the reduction in DSBs following TOP1 KD and RNase H1 OE is attributed to a decrease in transcriptional R-loops and trapped TOP1cc. Moreover, our findings suggest that transcriptional R-loops contribute to the trapping and stabilization of TOP1cc at highly expressed genes, as also suggested by the observed high enrichment of R-loops at genes with high TOP1cc (Figure 6I and J).

### Transcriptional DSBs as a driving force to cancer development

Endogenous DNA damage serves as a significant contributor to genomic instability in cancer (Tubbs and Nussenzweig, 2017). In line with this notion, we embarked on investigating the impact of physiological DSBs on the molecular alterations associated with the initiation and progression of breast cancer. Specifically, we focused on elucidating the role of transcriptional DSBs in this context. To address these questions comprehensively, we performed a thorough analysis that integrated mutational data from The Cancer Genome Atlas MC3 project (Ellrott et al., 2018), encompassing single nucleotide variant (SNV) mutational calls from breast cancer patients, with our own data obtained through sBLISS analysis of the breast cancer cell line MCF-7. Through an examination of break density, expression levels, TOP1 and R-loop densities, we aimed to discern potential disparities in transcriptional DSB accumulation between frequently mutated genes and the entire gene set. Our analysis as depicted in Supplementary Figure 8A-D, indicated no significant distinctions between frequently mutated genes and the broader gene population.

The prevailing understanding is that the majority of mutations in cancer are passengers and do not actively contribute to the initiation or progression of carcinogenesis (Martínez-Jiménez et al., 2020). To delve deeper into the involvement of physiological transcriptional DSBs in the early mutational events driving cancer initiation, we conducted a more focused analysis, specifically examining breast cancer driver genes(Nik-Zainal et al., 2016) in relation to non-functional highly mutated passenger genes. Remarkably, while passenger mutations were more prevalent across patients (Figure 7A), driver genes displayed a significantly higher density of breaks (Figure 7B). Intriguingly, driver genes also exhibited elevated expression levels and increased levels of TOP1 and R-loops (Figure 7C-E). This pattern persisted when comparing driver genes to the broader gene population (Supplementary Figure 8E-I). Notably, the expression levels of driver genes demonstrated a positive correlation with their break density, unlike passenger genes (Figure 7F and J). Both classes of mutated genes exhibited correlations between DSBs and R-loops (Supplementary Figure 8J and K), as well as between DSBs and TOP1 (Supplementary Figure 8L and M). Furthermore, a significant reduction in DSBs was observed in most driver genes following knockdown of TOP1 and/or overexpression of RNase H1 (Figure 7G). Upon further analysis based on expression levels, a clear trend emerged: highly expressed driver genes exhibited decreased DSBs in response to these manipulations, while lowly expressed driver genes did not show a similar pattern (Figure 7H and I). These findings strongly suggest an enrichment of transcriptional DSBs specifically at highly expressed driver genes. In contrast, passenger genes did not exhibit a significant decrease in DSBs after the manipulations (Figure 7K), implying an absence of transcriptional DSB enrichment in these genes.

**Figure 7:**
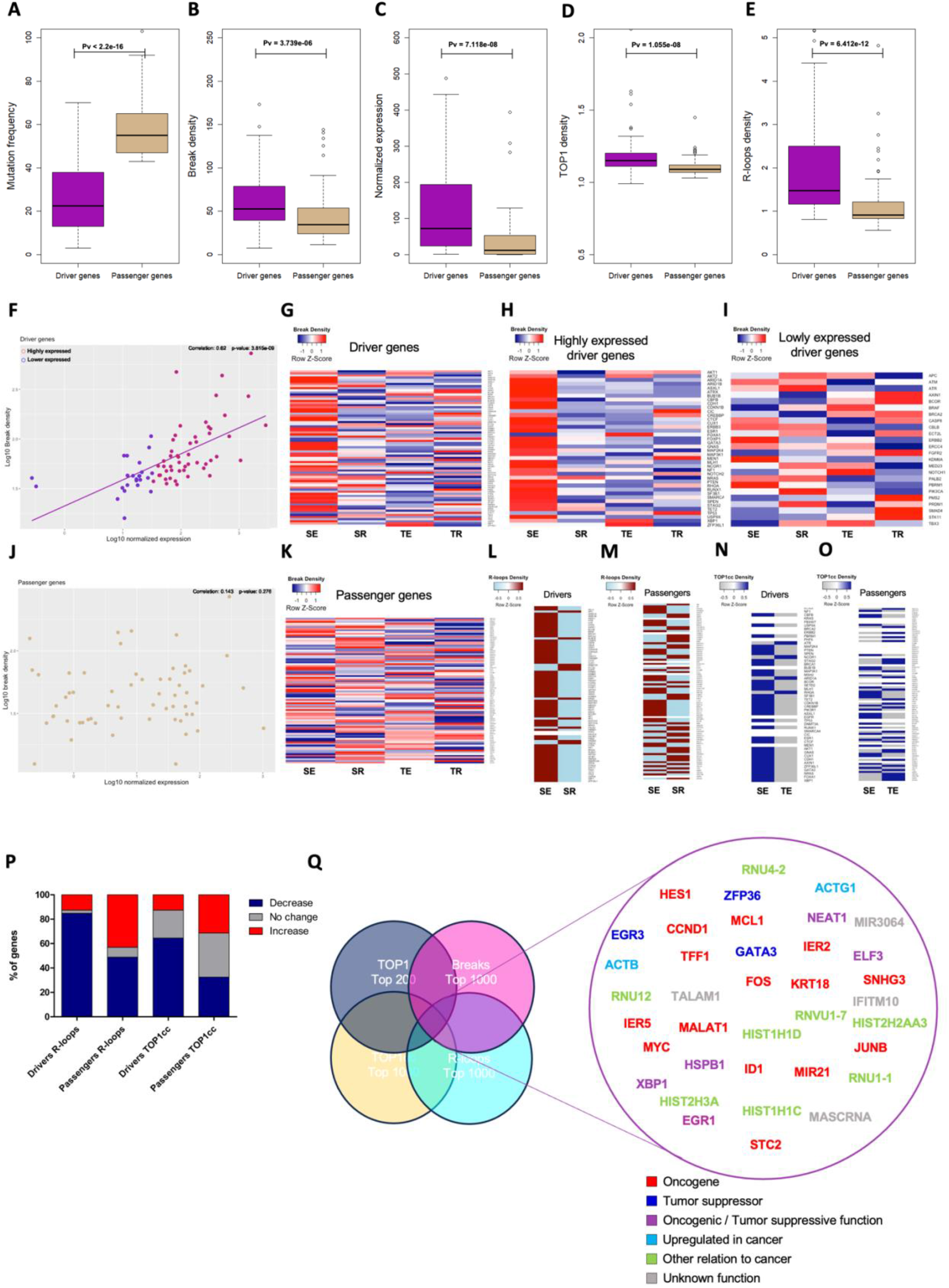
Cancer genes are enriched with transcriptional DSBs. **A.** Boxplot of mutation frequency at driver genes and frequently mutated passenger genes. **B.** Boxplot of break density at driver genes and frequently mutated passenger genes. **C.** Boxplot of normalized expression at driver genes and frequently mutated passenger genes. **D.** Boxplot of TOP1 density at driver genes and frequently mutated passenger genes. **E.** Boxplot of R-loops density at driver genes and frequently mutated passenger genes. **F.** Log-scaled scatter plot demonstrating the correlation between break density and expression levels at driver genes. Red circles are identifying highly expressed drivers used in the heatmap in **H**, blue circles are identifying lowly expressed drivers used in the heatmap in **I**. **G.** Color coded Heatmap of break density at driver genes with the different manipulations. **H.** Color coded Heatmap of break density at highly expressed driver genes with the different manipulations **I.** Color coded Heatmap of break density at lowly expressed driver genes with the different manipulations **J.** Log-scaled scatter plot demonstrating the correlation between break density and expression levels at passenger genes. **K.** Color coded Heatmap of break density at passenger genes with the different manipulations. **L.** Color coded heatmap demonstrating the direction of change of R-loops enrichment at driver genes after RNase H OE. **M.** Color coded heatmap demonstrating the direction of change of R-loops enrichment at passenger genes after RNase H OE. **N.** Color coded heatmap demonstrating the direction of change of TOP1cc enrichment at driver genes after TOP1 KD. **O.** Color coded heatmap demonstrating the direction of change of TOP1cc enrichment at passenger genes after TOP1 KD. **P.** Stacked bar graph demonstrating percentages of genes that decreased, didn’t change, or increased in the heatmaps mentioned earlier (L-O). **Q.** Genes that overlap the top lists of TOP1, breaks, TOP1cc, and R-loops with their available classification obtained from the literature. Gene enrichments are statistically significant and as follows, oncogenes PV= 1.103765e-27, tumor suppressors + dual function = 6.100708e-05, Actins = 2.772229e-05, Histones = 3.672962e-06.

To further validate these observations, we assessed the changes in R-loops and TOP1cc density upon overexpression of RNase H1 and knockdown of TOP1, respectively. Strikingly, while DSBs correlated with TOP1 and R-loops for both driver and passenger genes (Supplementary Figure 8 J-M), and there was no significant difference in TOP1cc density between drivers and passengers (Supplementary Figure 8N), only driver genes exhibited a decrease in R-loops upon RNase H1 overexpression and a decrease in TOP1cc density upon TOP1 knockdown (Figure 7L-P). These findings further imply that driver genes, but not passenger genes, are enriched with transcriptional DSBs mediated by TOP1cc trapping and R-loops. These results underscore the notion that driver mutations not only differ from passenger mutations in their impact on cancer fitness but also in their etiology.

Notably, when comparing driver genes to other similarly expressed genes, there is no significant difference in break density or R-loops density (Supplementary Figure 9A, B and C). This finding suggests that the disparities observed between driver and passenger genes are attributed to variations in their expression levels, rendering them susceptible to transcriptional DSBs.

Intriguingly, to determine if the observed differences between driver and passenger genes persist in cells before cancer transformation, we extended our investigation beyond the breast cancer cell line MCF-7. We sought to explore whether these patterns were evident in mammary epithelial cells, representing a pre-cancerous state. For this purpose, we conducted sBLISS analysis on the mammary epithelial cell line (HMLE) and simultaneously analyzed published RNA-seq data (GSM6774900). Astonishingly, we observed a similar pattern, reinforcing the presence of elevated expression and increased break density at driver genes compared to passenger genes (Supplementary Figure 9D and E).

Furthermore, we harnessed the breast cancer progression model, utilizing the normal MCF-10A cells and their malignant counterpart harboring RAS mutation (RAS-mutated MCF-10A cell line) (Salah, Itzhaki and Aqeilan, 2014). Through sBLISS experiments and the examination of published RNA-seq data (Tracy et al., 2018), we consistently observed that driver genes exhibited higher expression levels and displayed greater break density than passenger genes in both non-malignant and RAS-transformed MCF-10A cells (Supplementary Figure 9F-I). Interestingly, we found no significant difference in the expression or break density of driver genes between the normal and the transformed cell line (Supplementary Figure 9J and K), indicating that the enrichment of transcriptional DSBs at driver genes is indeed present even before cancer transformation events occur. These findings suggest that the unique transcriptional DSB landscape associated with driver genes is a feature preserved throughout various cellular stages, including pre- cancerous and cancerous states.

To uncover the distinct characteristics of genes acting as hotspots for transcriptional DSBs, we employed a different approach. By identifying overlapping genes from the top lists of DSBs obtained through sBLISS, TOP1 and TOP1cc from ChIP-seq, and R-loops from DRIP-seq, all performed on non-treated MCF-7 cells, we obtained a comprehensive understanding (Figure 7Q, Supplementary Figure 9L-O). Remarkably, out of the 37 genes that intersected across all four lists, 33 genes were implicated in cancer initiation and/or progression. This collection included 15 oncogenes, such as *MYC*, *FOS*, *HES1*, *JUNB*, and *CCND1*. Additionally, three genes were classified as tumor suppressors (*EGR3*, *ZFP36*, and *GATA3*), while five genes demonstrated dual roles as either oncogenes or tumor suppressors depending on the specific context (*XBP1*, *EGR1*, *HSPB1*, *NEAT1*, and *ELF3*). Notably, two genes in the list belonged to the actin family, namely *ACTB* and *ACTG1*, and their aberrant expression and dysregulation have been observed in various types of cancer, contributing to tumor progression and metastasis (Guo et al., 2013; Gu et al., 2021; Suresh and Diaz, 2021). The remaining genes in the list have limited information regarding their role in cancer; however, they belong to gene families known to play critical roles in cancer biology. Specifically, four genes belong to the histone family (*HIST1H1D* (Huang et al., 2021), *HIST1H1C*, *HIST2H3A*, and *HIST2H2AA3* (Monteiro et al., 2014) and mutations in histone genes have been associated with cancer (Funato and Tabar, 2018; Soshnev et al., 2021). Furthermore, four genes belong to the small nuclear RNAs (snRNAs) family, specifically *RNU1*-1, *RNU1*-7, *RNU4*-2, and *RNU12*. While these particular snRNAs may not commonly be associated with cancer, dysregulation or alterations in the broader snRNA family and splicing machinery have implications for cancer development and progression (Suzuki et al., 2019; Zhang et al., 2021).

Taken together, these findings provide substantial evidence that transcriptional DSBs, facilitated by TOP1cc trapping and R-loops, are intimately associated with early molecular changes in breast cancer. Moreover, the disparities between driver mutations and passenger mutations extend beyond their molecular functions and contributions to cancer fitness; they also encompass differences in their etiology. The higher expression levels, enrichment of R-loops, and elevated presence of TOP1 in driver genes suggest that transcriptional DSBs could be the underlying driving force behind mutagenesis in cancer.

## Discussion

In this study, we conducted a comprehensive investigation to elucidate the mechanisms and consequences of transcriptional DSBs in the context of cancer development. Our analyses revealed a strong association between DSBs and highly transcribed genes, with elevated break density observed at active promoters, enhancers, and transcriptional transition regions. Moreover, we demonstrated that these transcriptional DSBs are influenced by the interplay between TOP1 and R-loops. Specifically, highly expressed genes exhibited a higher density of breaks, correlated with increased TOP1 levels and R-loop formation. Depletion of TOP1 and R-loops resulted in a significant reduction in DSBs at these highly transcribed genes, underscoring their crucial roles in transcription-associated genomic instability. Furthermore, we investigated the contribution of transcriptional DSBs to cancer initiation and progression. Driver genes, implicated in cancer fitness, displayed higher break density, increased expression levels, and elevated levels of TOP1 and R-loops compared to passenger genes. Manipulations of TOP1 and R-loops selectively decreased DSBs in highly expressed driver genes, providing evidence for their enrichment in transcriptional DSBs and distinct etiology. These findings deepen our understanding of driver mutations by elucidating the role of physiological DSBs as an additional factor facilitating their occurrence alongside Darwinian evolution. By unraveling the molecular characteristics underlying these driver mutations, our research provides valuable insights into the complex interplay between genomic instability, transcriptional processes, and cancer development and highlights the regulatory roles of TOP1 and R-loops in governing DSBs within hypertranscribed genes implicated in carcinogenesis.

### Transcriptional DSBs at TSS are different from gene body

Our findings propose an intriguing model highlighting the distinct production of transcriptional DSBs at transcription start sites compared to those within the gene body. Specifically, we observed that depletion of TOP1 and reduction of R-loops specifically resulted in reduced DSBs at the gene bodies of highly expressed genes, while leaving the DSBs at TSS relatively unaffected. This observation is consistent with the mapping of TOP1 cleavage complexes along the gene body, which showed depletion of TOP1cc at promoters compared to gene body (Figure 6D). Moreover, this finding aligns with the recent discovery that DSBs at TSS are primarily produced by the high-mobility group AT-hook 2 protein (HMGA2), a key factor required for transcription initiation (Dobersch et al., 2021). Our previous study also demonstrated differential patterns of DSBs at regulatory elements such as promoters, strong enhancers, and insulators (Hazan et al., 2019). Furthermore, our current investigation revealed a slight increase in break density at promoters and enhancers following TOP1 and R-loop depletion. This observation suggests that TOP1 and R-loops may serve distinct functions at these regulatory regions beyond their association with hypertranscription. This supposition aligns cohesively with the findings of a recent study by (Ray et al., 2022), which underscored the distinct susceptibility of promoters to specific forms of DNA damage. This alignment fortifies our perspective on the distinct origins of DNA damage between TSS and gene bodies, potentially indicating that TOP1 and R-loops could act as protective agents at promoters, safeguarding against these threats.

Together, our data support a model in which transcriptional DSBs at TSS are produced differently from those within the gene body and that TOP1 and R-loops cause DSBs at gene bodies, thereby establishing them as distinctive attributes of transcriptional DSBs.

### The Connection between R-loops and TOP1

Remarkably, our results consistently demonstrate that the depletion of TOP1 and R-loops elicits similar effect, and the combined manipulation fails to produce a synergistic effect. This observation suggests a coordinated process in which the presence of both factors is necessary to initiate the generation of DSBs through a shared pathway, potentially involving the dual processing of TOP1 and R-loops (Cristini et al., 2019). In this scenario, a SSB arises from TOP1 activity, while another SSB results from an R-loop, ultimately leading to the formation of DSBs. Our data strongly support this mechanistic model and indicate that it holds true even under physiological conditions. Indeed, the unexpected effect of both manipulations on expression levels, suggests the involvement of complex compensatory mechanisms. It is plausible that the compensatory responses triggered by each manipulation individually may interact in a way that interferes with each other when both manipulations are present simultaneously. This interference could lead to additional stress on the transcriptional machinery, resulting in the observed decrease in expression at highly transcribed genes.

Consistent with the literature, our data show that TOP1 knockdown increases the presence of R-loops at highly transcribed genes. However, this increase in R-loops does not correspond to a concurrent increase in DSBs, further supporting that both TOP1 and R-loops are essential components for DSB formation. Furthermore, our data provide an additional angle to the intricate mechanism whereby R-loops facilitate the trapping of TOP1cc and subsequent DSB formation.

### Estrogen responsiveness and DSBs

Previously, R-loops have been shown to be the main cause of estrogen-induced DNA damage (Stork et al., 2016), pointing towards a role of R-loops in transcriptional DSBs. Our sBLISS data on E2 responsive genes after TOP1 KD or RNase H1 OE shows induction of breaks at these genes after estradiol treatment, further validating a causing effect of transcription on DSBs formation. Furthermore, TOP1 KD or RNase H1 OE significantly attenuate the effect of estradiol, validating the involvement of TOP1 and R-loops in transcriptional DSBs formation.

### Increased expression upon TOP1 KD and RNase H1 OE

Our findings reveal a noteworthy correlation between decreased DSBs and enhanced expression levels of highly transcribed genes following TOP1 knockdown and RNase H1 OE. This observation aligns with the prevailing concept that DSBs act as impediments to the progression of the transcriptional machinery. Interestingly, despite the importance of TOP1 activity in facilitating transcription, our results indicate that the remaining 30% of the protein is sufficient for efficient transcription, implying that TOP1 is often overexpressed in cancer cells beyond what is strictly necessary for cellular processes.

Furthermore, the reduction in TOP1cc resulting from TOP1 KD and RNase H1 OE provides a mechanistic explanation for the observed increase in gene expression. It is well-documented that TOP1cc represents transcription-blocking lesions (Cristini et al., 2016), and thus, the decrease in TOP1cc levels can alleviate the transcriptional impediments, consequently promoting gene expression.

Collectively, our data support the notion that decreased DSBs, brought about by TOP1 KD and RNase H1 OE are associated with elevated expression of highly transcribed genes. This sheds light on the intricate relationship between DSB formation, TOP1 activity, and transcriptional regulation, underscoring the potential implications of these mechanisms in cancer biology.

### Transcriptional DSBs and mutations

The findings of our study shed light on the significance of transcriptional DSBs as a driving force in the development and progression of breast cancer. Our analysis revealed that driver genes exhibited significantly higher break density compared to both passenger genes and the overall gene population. Furthermore, the enrichment of R-loops and TOP1, along with the decreased DSBs following manipulations of TOP1 and RNase H1, in driver genes supports the notion that transcriptional DSBs facilitated by TOP1cc trapping and R-loops contribute to the underlying mechanisms of mutagenesis in breast cancer. Importantly, our data challenge the existing knowledge regarding the formation of driver mutations, as we demonstrate that transcriptional DSBs play a significant role beyond random chance. This novel insight proposes an update to our understanding of the mechanisms governing driver mutation formation. Indeed, it is essential to clarify that the enrichment of transcriptional DSBs at driver genes is likely a consequence of their increased transcriptional levels and not an independent or intrinsic feature unique to driver genes. Our data strongly support the notion that the presence of transcriptional DSBs at driver genes can be attributed to their higher levels of transcription, which in turn make them more prone to the formation of DSBs. Conversely, the absence of transcriptional DSBs at passenger genes can be explained by their lower expression levels, rendering them less susceptible to DSB formation.

It is important to note that when comparing driver genes with other similarly expressed genes, there is no significant difference in the occurrence of transcriptional DSBs between drivers and these similarly expressed non-functional passenger genes. This reinforces the link between transcriptional activity and the enrichment of DSBs at driver genes. The presence of DSBs at driver genes is likely driven by their active transcriptional state, as highly expressed genes are inherently more vulnerable to DNA damage due to their increased transcriptional activity. Conversely, passenger genes, which lack significant functional roles in cancer initiation and progression, tend to be located within genes that are lowly expressed. As a result, they are depleted of transcriptional DSBs, highlighting the importance of transcription in driving DSB formation and underscoring the significance of active transcription in this process.

Furthermore, our investigation also identified several genes intersecting the lists of DSBs, TOP1, TOP1cc, and R-loops, with a majority of them being linked to cancer initiation and/or progression. These findings highlight the association of transcriptional DSBs with well-established oncogenes, tumor suppressors, and genes from critical families such as histones and snRNAs. While further research is needed to unravel the exact mechanisms and downstream consequences of transcriptional DSBs, our study provides compelling evidence for their involvement in early molecular changes in breast cancer. Understanding the etiology and molecular characteristics of driver mutations, including their relationship with transcriptional DSBs, not only enhances our knowledge of cancer biology but also holds potential for the development of targeted therapeutic strategies.

## Acknowledgments

We thank Ameen Haj-Yahia for his assistance in ChIP-seq. We are grateful to all the Aqeilan lab members for fruitful discussion and advise. This study was funded by grants from the European Research Council (ERC) [No. 682118] and Israel Science Foundation (ISF) [No. 1056/21].

## Method details

### Cell culture

MCF7 (HTB-22) were grown in RPMI supplemented with 10% (vol/vol) FBS (GIBCO), glutamine, and penicillin/streptomycin (Beit-Haemek). HMLE cells were grown in Promocell mammary epithelial cell basal media (C-21010) with added supplements (c- 93110) whereas MCF10A were grown on DMEM/F12 supplemented with 5% Horse serum, 20 ng/ml EGF, 0.5 mg/ml Hydrocortisone, 100 ng/ml Cholera toxin, 10 mg/ml Insulin and Pen/Strep. MCF-7 cells were synchronized in the G1 cell cycle stage by culturing in serum-free media for 72 hours followed by replacing the media with 10% FBS media for 5 hours. Cells were grown at 37°C under a humidified atmosphere with 5% CO2. Cells were routinely authenticated by STR profiling, tested for mycoplasma, and cell aliquots from early passages were used.

### Transient transfection and plasmids

Transient transfection of siTOP1 (Dharmacon; SMARTpool siRNA, L-005278-00-0005) and siSc (Dharmacon; D-00181010) was achieved using lipofectamine (ThermoFisher). Cells were seeded to reach a confluency of 60-70% on the day of transfection. Transfection solution was prepared by mixing 1.5ml serum/antibiotic-free RPMI with 30μL lipofectamine and incubating for 5 minutes, before adding a mixture of 1.5ml RPMI and 0.6nmoles of siRNA followed by 20 minutes incubation at RT. The mixture was then added to the cells in a 10cm plate cultured in antibiotic/serum-free media. The media was replaced by fully supplemented media after 5-6 hours, and cells were incubated in the incubator for 48 hours.

For RNase H1 overexpression, RNase H1 plasmid and its empty control plasmid containing only GFP, a kind gift from Dr. Sara Selig (Molecular Medicine Laboratory, Rambam Health Care Campus and Rappaport Faculty of Medicine, Technion, Haifa 31096, Israel), were used. GFP expression was used as an indicative of RNase H1 overexpression. The plasmid was prepared as described in (Sagie et al., 2017).

The knockdown in TOP1cc ChIP-seq and DRIP-seq experiments were done using the doxycycline inducible shTOP1 plasmid that was kindly obtained from Yilun Liu.

### Immunofluorescence assay

Cells were seeded on coverslips to a 50-70% confluency at the time of fixation. Fixation was done using 4% paraformaldehyde for 10 minutes, followed by three washes with 1X PBS. Cells were then permeabilized with 0.25% Triton X for 7 minutes followed by another three washes with 1X PBS. This was followed by blocking with 10% goat serum for 1 hour. Cells were then stained with anti-γ-H2AX antibody (abcam, ab26350) using 1:1000 dilution, and anti-53BP1 anti-body (abcam, ab36823) using 1:500 dilution, and incubated in a humidity chamber in the cold room overnight. After three washes with PBS, cells were incubated with the secondary antibodies in a humidity chamber for one hour at room temperature, and then stained with Hoechst stain (1:2000 dilution in 1X PBS) followed by two washes with PBS. Coverslips were then mounted, and immunofluorescent pictures were taken using confocal microscopy. For TOP1cc (MABE1048), the same protocol was used with the addition of treating the permeabilized cells with 0.1% SDS for 5 minutes before the blocking to render the covalent bond accessible for the antibody.

### CEL-seq

Total RNA was extracted using TRI reagent (BioLabs) according to the manufacturer specifications. RNA concentration was measured using Qubit Flex Fluorometer (ThermoFisher Scientific). RNA sequencing libraries were prepared using the CEL-Seq2 protocol in the Technion genomic Center (TGC), as published by (Hashimshony et al., 2012), with one modification; instead of single-cells as input, 2 ng purified RNA was taken as input for library preparation. The CEL-Seq2 libraries were analyzed for average fragment size using Agilent 2200 TapeStation (Agilent) and concentration was measured using Qubit Flex Fluorometer (ThermoFisher Scientific). The libraries were sequenced on the Illumina NextSeq 2000 sequencer (Illumina), 12 bases for read 1 and 88 bases for read 2. Demultiplexing was performed in two steps. First, Illumina demultiplexing was performed using bcl2fastq Illumina software with the following parameters: barcode-mismatches =1, minimum-trimmed-read-length = 0, and mask-short-adapter-reads = 0. Second, Cell-seq demultiplexing using the pipeline described in (Hashimshony et al., 2012) was executed with the following parameters: min_bc_quality = 10, bc_length = 6, umi_length = 6, and cut_length = 88. RNA measurements, library preparation and sequencing were performed by the Technion Genomics Center, Technion, Israel.

### Western blot

Protein lysates were prepared by incubating cells in lysis buffer containing 50 mM Tris (pH7.5), 150 mM NaCl, 10% glycerol, 0.5% Nonidet P-40 (NP-40), with protease and phosphatase inhibitors (1:100). Samples were run on SDS-PAGE gel for 90-120 minutes and then blotted on nitrocellulose membrane using semi-dry blotting machine (Biorad). Two antibodies were used, Topoisomerase 1 antibody (Cat# ab3825) for validation of siRNA knockdown, and GAPDH antibody (Calbiochem; CB1001), as a housekeeping gene antibody.

### Transcription induction using Estradiol

MCF-7 cells were hormonally starved by growing in RPMI media supplemented with 10% charcoal stripped fetal bovine serum for 48 hours. Transcription was induced by replacing the media with RPMI media supplemented with charcoal stripped FBS and 100nM β- Estradiol (Sigma-Aldrich) (pre-dissolved in ethanol) and incubated for 1 hour.

### In-suspension break labelling *in situ* and sequencing (sBLISS)

sBLISS was performed as described in (Bouwman et al., 2020). Briefly, 10^6^ cells were fixed in 2% paraformaldhyde in 10%FCS/PBS for 10min at room temperature. Fixation was quenched with 125 mM glycine for 5min at RT, and another 5min on ice, followed by two washes in ice cold PBS. The cells were lysed for 60min on ice and the nuclei were permeabilized for 60min at 37°c. Then, nuclei were washed twice with CutSmart Buffer supplemented with 0.1% Triton X-100 (CS/TX100), and DSB ends were in situ blunted with NEB’s Quick Blunting Kit for 60min at RT. Blunted nuclei were washed twice with 1x CS/TX100 before proceeding with in situ ligation of sBLISS adapters to the blunted DSB ends. Adaptor ligation was performed with T4 DNA Ligase for 20-24h at 16C and supplemented with BSA and ATP. After ligation, nuclei were washed twice with 1x CS/TX100 and genomic DNA was extracted with Proteinase K at 55°c for 14-18h while shaking at 800rpm. Afterward, Proteinase K was heat-inactivated for 10 min at 95°c, followed by extraction using Phenol:Chloroform:Isoamyl Alcohol, Chloroform, and ethanol precipitation. The purified DNA was sonicated in 100 μL ultra-pure water using Covaris M220 for 60s. Sonicated samples were concentrated with AMPure XP beads (Beckman Coulter) and fragment sizes were assessed using a BioAnalyzer 2100 (Agilent Technologies) to range from 300bp to 800bp with a peak around 400-600bp. The sonicated DNA was in vitro transcribed using MEGAscript T7 Kit for 14 h at 37°c. After RNA purification and ligation of the 3^’^-Illumina adaptor, the RNA was reverse transcribed. The final step of library indexing and amplification was performed using NEBNext® Ultra™ II Q5® Master Mix.

### Chromatin immunoprecipitation sequencing (ChIP-seq)

MCF7 cells (∼10^7^) were crosslinked with 1% formaldehyde (methanol free, Thermo Scientific 28906) for 10 min at room temperature and quenched with glycine, 125 mM final concentration. Fixed cells were washed twice in PBS and incubated in lysis buffer (10mM EDTA, 0.5% SDS, 50mM Tris-HCl pH=8, and protease and phosphatase inhibitors) for 30min on ice. Cells were sonicated using bioruptor sonicator to produce chromatin fragments of ∼200-300 bp. The sheared chromatin was centrifuged 10min at maximum speed. From the supernatant, 2.5% were saved as input DNA and the rest was diluted in dilution buffer (50mM TRIS-HCl pH8, 0.01%SDS, 150mM NaCl, and 1% Triton X-100). The chromatin was immunoprecipitated by incubation with 5 μl of anti-TOP1cc antibody (MABE1048). Immune complexes were captured with protein G Dynabeads. Immunoprecipitates were washed once with low salt, twice with high salt buffer, and twice with LiCl buffer, and twice with TE buffer. The chromatin was eluted from the beads with 300 μl of elution buffer (100mM sodium bicarbonate and 1% SDS) and incubated overnight at 65°C to reverse the cross-linking. Samples were treated with proteinase K at 45 °C for 2 h. DNA was precipitated by phenol/chloroform/isoamylalcohol extraction. The ChIPed and the Input DNA were used to prepare libraries by Kappa Hyperprep kit and sequenced in Nextseq (illumine).

### DNA-RNA immunoprecipitation (DRIP-seq)

DRIP-seq was done according to the published protocol (Sanz and Chédin, 2019).

### Bioinformatic analysis and data processing sBLISS analysis

sBLISS fastq files are first de-multiplexed by sample barcodes. Quality control is applied with trim_galore to remove residual adapters, trim reads to base quality of at least 20 and filter out short reads with size smaller than 20 bp. Initial and final sample qualities are evaluated with fastqc. Quality processed fastq files are aligned to GRCh38 assembly with hisat2 and then sorted and indexed with samtools. Resulting bam files are de-duplicated with umi-tools employing genomic coordinates and Unique Molecular Identifiers (UMI). Custom python and R scripts are applied to identify read start position and convert bam files to break bigwig format for downstream analysis. A custom R script is applied to discard a blacklist of positions (mainly within centromeres). For most analyses, breaks are aggregated over genomic tiles using R tileGenome utility.

### CEL-Seq analysis

Quality control is applied on CEL-Seq fastq files with trim_galore to remove residual adapters, polyA tails, trim reads to base quality of at least 20 and filter out short reads with size smaller than 20 bp. Initial and final sample qualities are evaluated with fastqc. Quality processed fastq files are then aligned to GRCh38 transcriptome using salmon aligner in it mapping mode while enabling the write_mapping key to create a sam file of transcripts. The sam file is deduplicated with a custom python script which scans for primary alignments only and deduplicates with respect to the transcript and Unique Molecular Identifiers (UMI). Non-duplicated primary alignments and all their associated secondary alignments are kept. Salmon aligner is executed again in its alignment mode, with sam deduplicated file as input resulting in a de-duplicated transcript count table.

### DRIP-Seq analysis

DRIP-Seq peak file is downloaded from GEO datasets - GSE81851. The peaks selected for analysis are for T0-Input after liftover to GRCh38.

### ChIP-Seq analysis

ChIPed and input fastq files are quality controlled with trim_galore to remove residual adapters, trim reads to base quality of at least 20 and filter out short reads with size smaller than 20 bp. Initial and final sample qualities are evaluated with fastqc. The quality processed fastq files are aligned to GRCh38 genome with hisat2 followed by sorting and indexing with samtools. The utility bamCompare of deepTools is applied on the bam files to create a peak bigwig file. This operation is set to ignore duplicates, discard a blacklist of regions, normalize by RPKM, and compare by ratio in tiles of 50 bp

### Algorithms used for plotting the data

Heatmaps were plotted using the following pipeline. First deepTools computeMatrix is applied with center reference-point. Next, an R script discards outlier spikes from the matrix. Finally, deepTools plotHeatmap is performed. In two-part plots, each part is defined as an independent region.

Regions of chromatin states are based on GSE57498 for HMEC cell line. For each of the regions, breaks are counted within that region and compared between treated samples or between treated sample and an expected count. The expected count is calculated as the proportion of breaks expected due to regions size relative to overall genome size. Plots are created with base R functions.

Figures showing break percentage at high expression/ R-loops/ TOP1 were generated using the following pipeline, Downsized breaks are counted across the gene body for all genes-of-interest (e.g., 1000 topmost expressed genes) and summed up for each of the treated samples. Downsizing is performed to reduce possible bias related to library preparation. An expected break value is also calculated based on the relative occupancy of the genes-of-interest relative to overall genome size. Bar-plots are created with base R functions.

For plots showing breaks distribution across gene body, each gene-of-interest is divided into 30 successive regions and breaks are counted separately in each region. In addition, for each gene, two flanks of size 3000 bp are subdivided into 10 regions and breaks are counted in the flanks. The plot shows break counts for several treated samples after normalization of each curve between 0 and 100.

For color coded heatmaps, Break density is assigned to each gene-of-interest (cancer related genes) and plotted as a heatmap after ranking by SE sample. Heatmap is created with heatmap.2 function of R gplots package.

Figures with bars and boxplots, and log-log scatter plots are created using base R functions.

## Data availability

Raw data are available upon request.

## Supplementary Figures

**Supplementary Figure 1:**
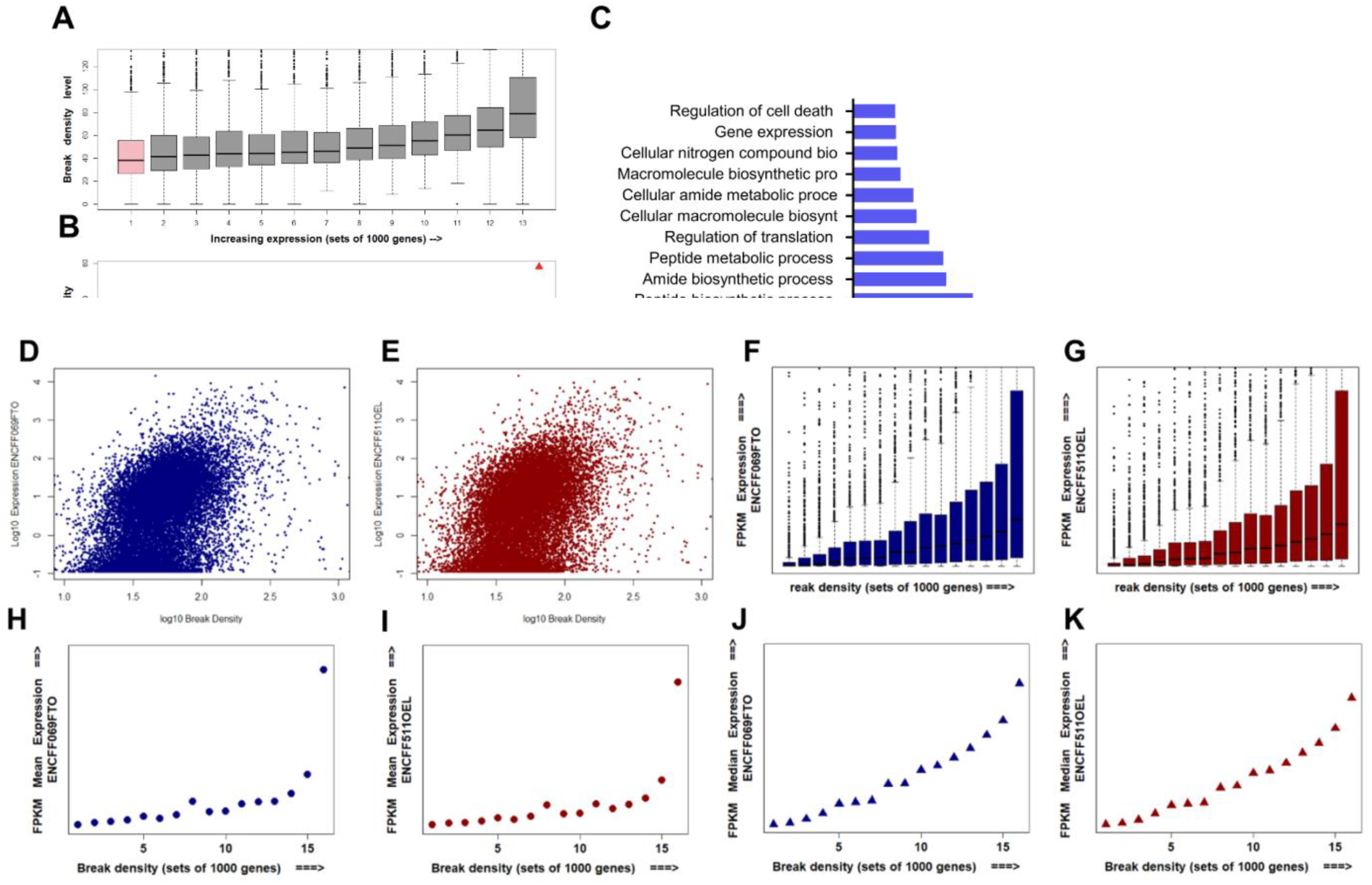
DSBs are enriched at highly expressed genes. **A**. Boxplot showing the correlation between break density and expression level, genes were grouped into 13 groups with increasing expression, the 13th group being the group with the highest mean expression. **B.** Median of break density positively correlates with break density. A zoomed log scaled scatter plot showing the correlation between expression levels and TOP1 level, each dot represents a gene. **C**. Gene ontology analysis of the 400 top broken genes in MCF-7 cells. **D** and **E**. A zoomed log scaled scatter plot showing a positive correlation between break density and expression levels; each dot represents a gene, and for each gene break density and expression were measured and normalized to gene size. **F** and **G**. Boxplot showing the correlation between break density and expression level. **H** and **I.** Mean expression (TPM) positively correlates with break density. **J** and **K.** Median expression (TPM) positively correlates with break density.

**Supplementary Figure 2:**
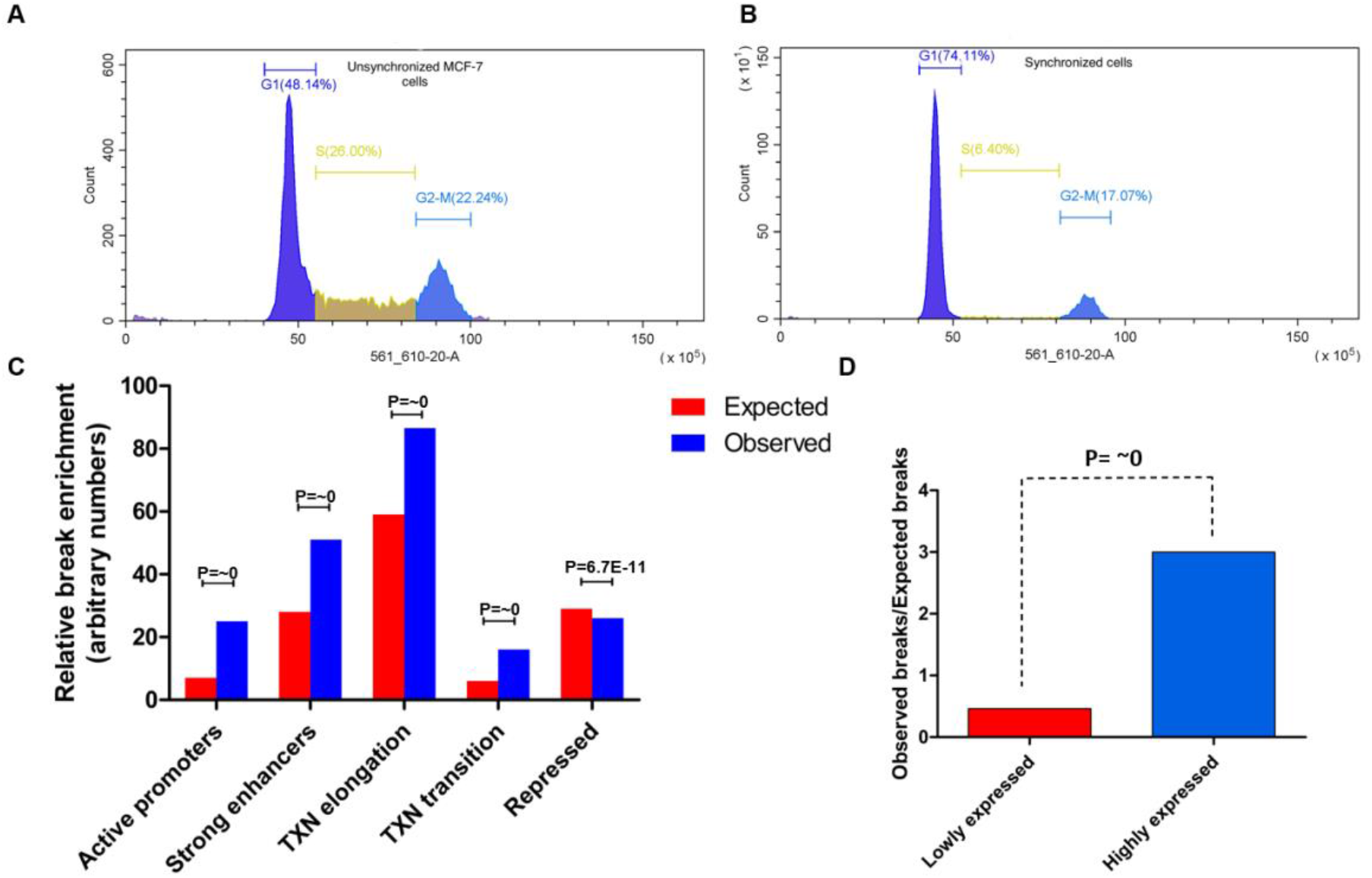
breaks are enriched at active regions in non-cycling MCF-7 cells. A-B. Cell cycle analysis for MCF-7 cells unsynchronized (**A**) and MCF-7 cells serum starved for 48 hours (**B**). **C.** The distribution of DSBs in synchronized breast cancer MCF7 cells along ChromHMM-defined chromatin states of HMEC, bar height is break enrichment relative to other chromatin states. **D.** Ratio of observed breaks vs expected breaks at highly and lowly expressed genes.

**Supplementary Figure 3:**
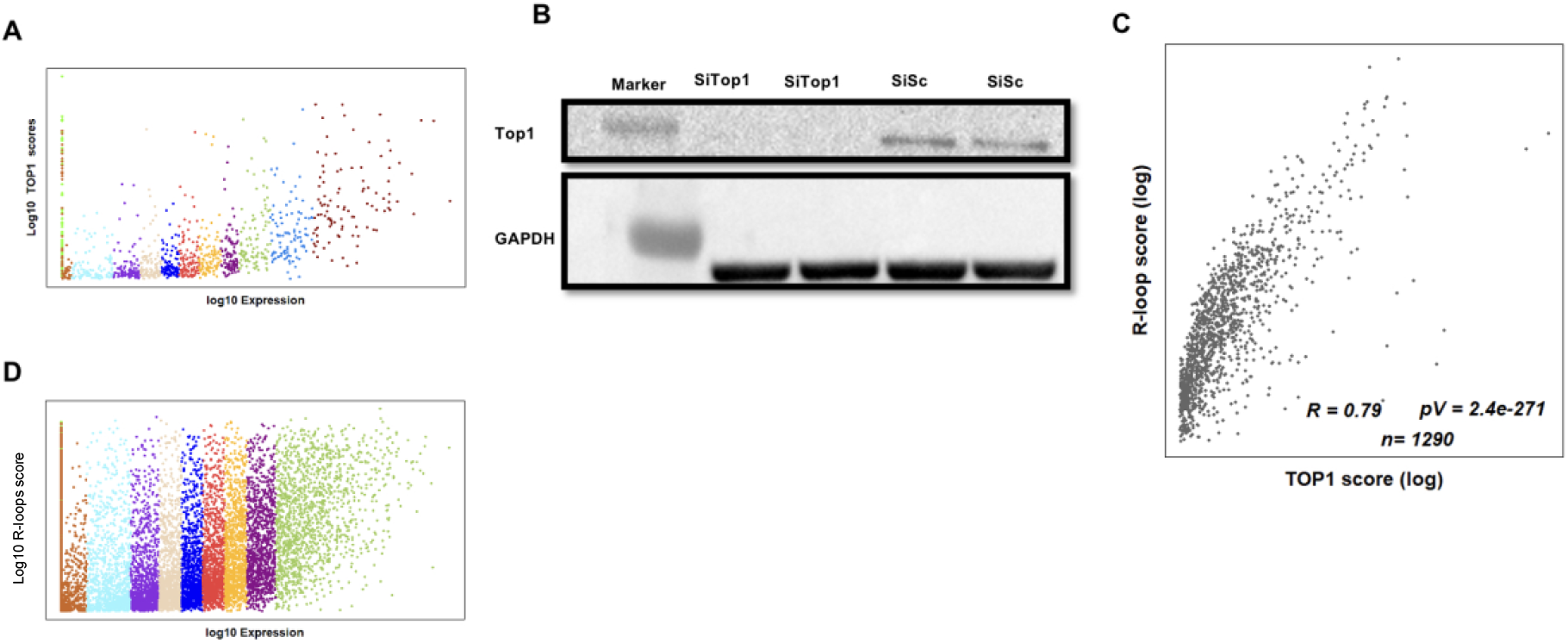
Transcriptional DSBs are associated with TOP1 and R-loops. **A.** A log scaled scatter plot showing positive correlation between expression and TOP1 levels; each dot represents a gene, and each color represents a gene group. **B.** Validation of TOP1 knockdown by Western Blot. **C**. A log scaled scatter plot showing the correlation between R loops and TOP1 levels, each dot represents a gene. DRIP data was extracted from GSE81851. **D.** A log scaled scatter plot showing positive correlation between expression and R-loop levels; each dot represents a gene, and each color represents a gene group.

**Supplementary Figure 4:**
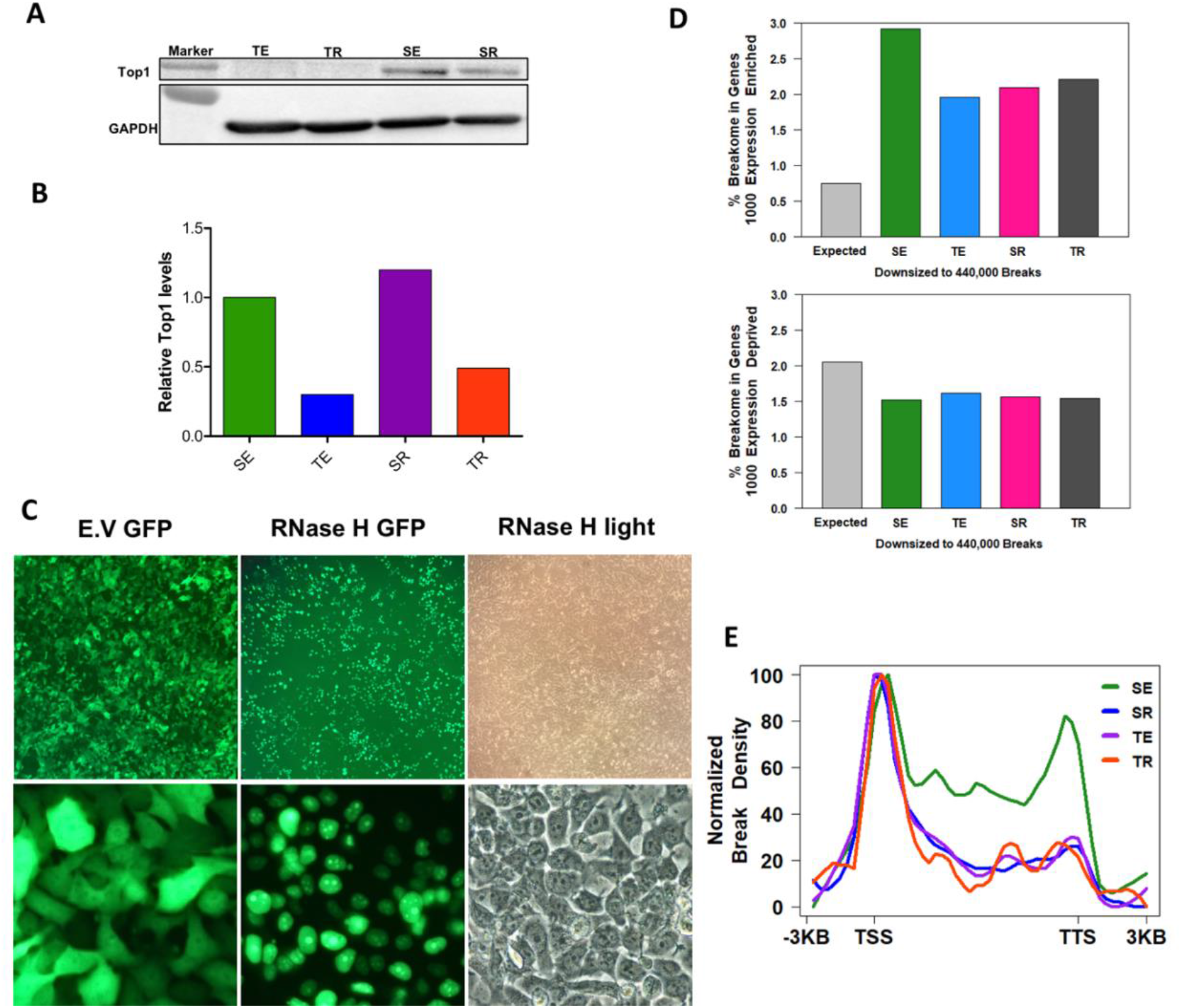
RNase H1 overexpression and TOP1 knockdown decrease break enrichment at gene body of highly expressed genes. **A**. Validation of TOP1 knockdown using western blotting. **B.** Quantification of relative TOP1 levels after TOP1 knockdown. **C.** GFP pictures for RNase H1- GFP overexpression. **D.** Breakome percentage in 1000 most transcribed genes (top) and 1000 least transcribed genes (bottom) with the different manipulations, expression data was extracted from encode projects ENCFF069FTO_FPKM and ENCFF511OEL_FPKM for MCF7. **E.** Plot showing break density at gene body of the 1000 highest transcribed genes with the different manipulations. P-values were calculated using chi-square test.

**Supplementary Figure 5:**
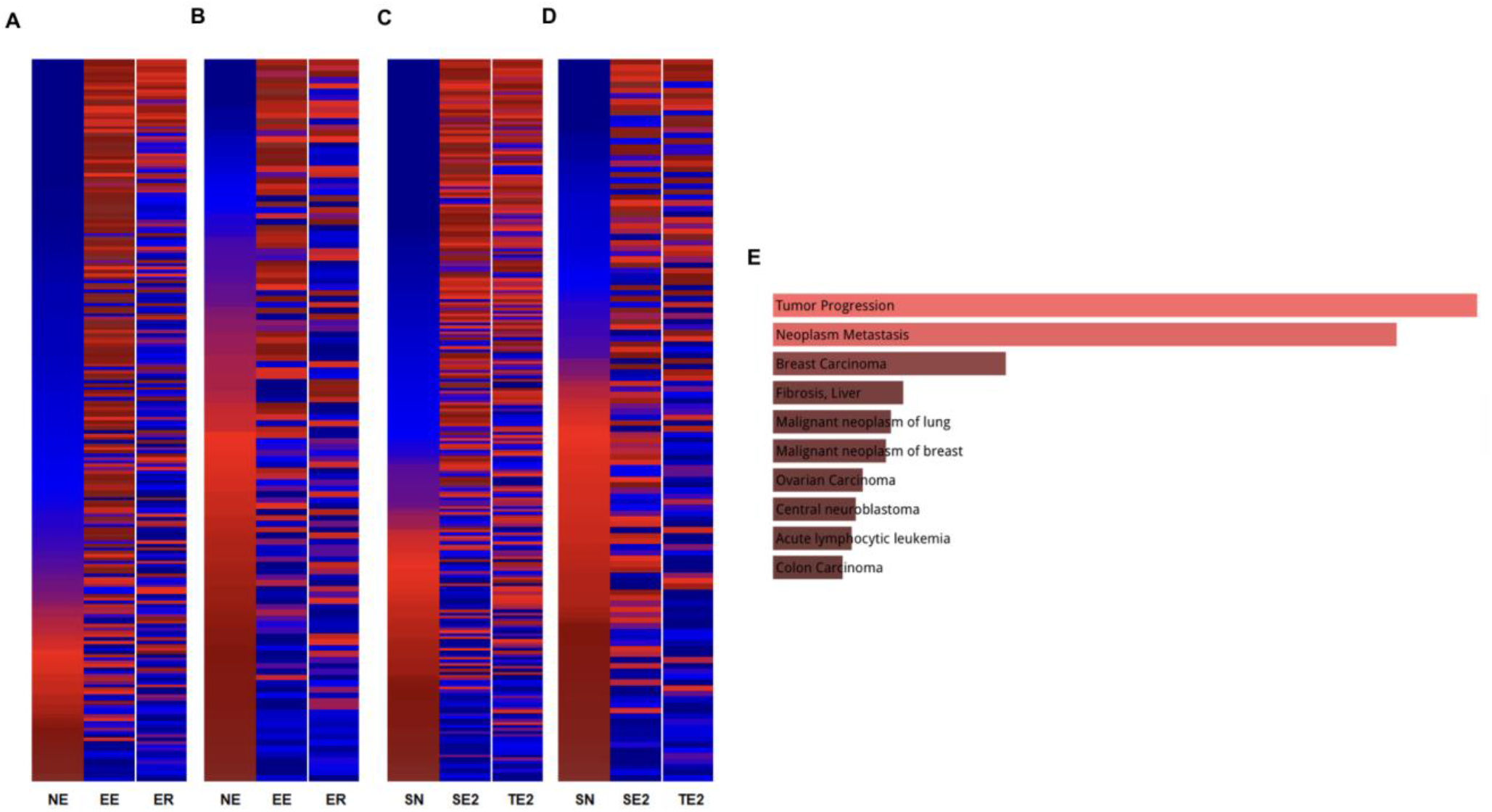
Estradiol associated DSBs are mediated by R-loops and TOP1. **A.** and **B.** Color coded heatmap showing the change in break density between control (NE), cells incubated with estradiol (EE), cells incubated with estradiol and overexpressing RNase H1 (ER), for estrogen positive responsive genes (E) and negative responsive genes (F). **C.** And **D.** Color coded heatmap showing the change in break density between control (SN), cells incubated with estradiol (SE2), cells knocked down for TOP1 (TN), for estrogen positive responsive genes (G) and estrogen negative responsive genes (H). **E.** Disease related pathways for estradiol responsive genes using DisGeNET database (Piñero et al., 2021).

**Supplementary Figure 6:**
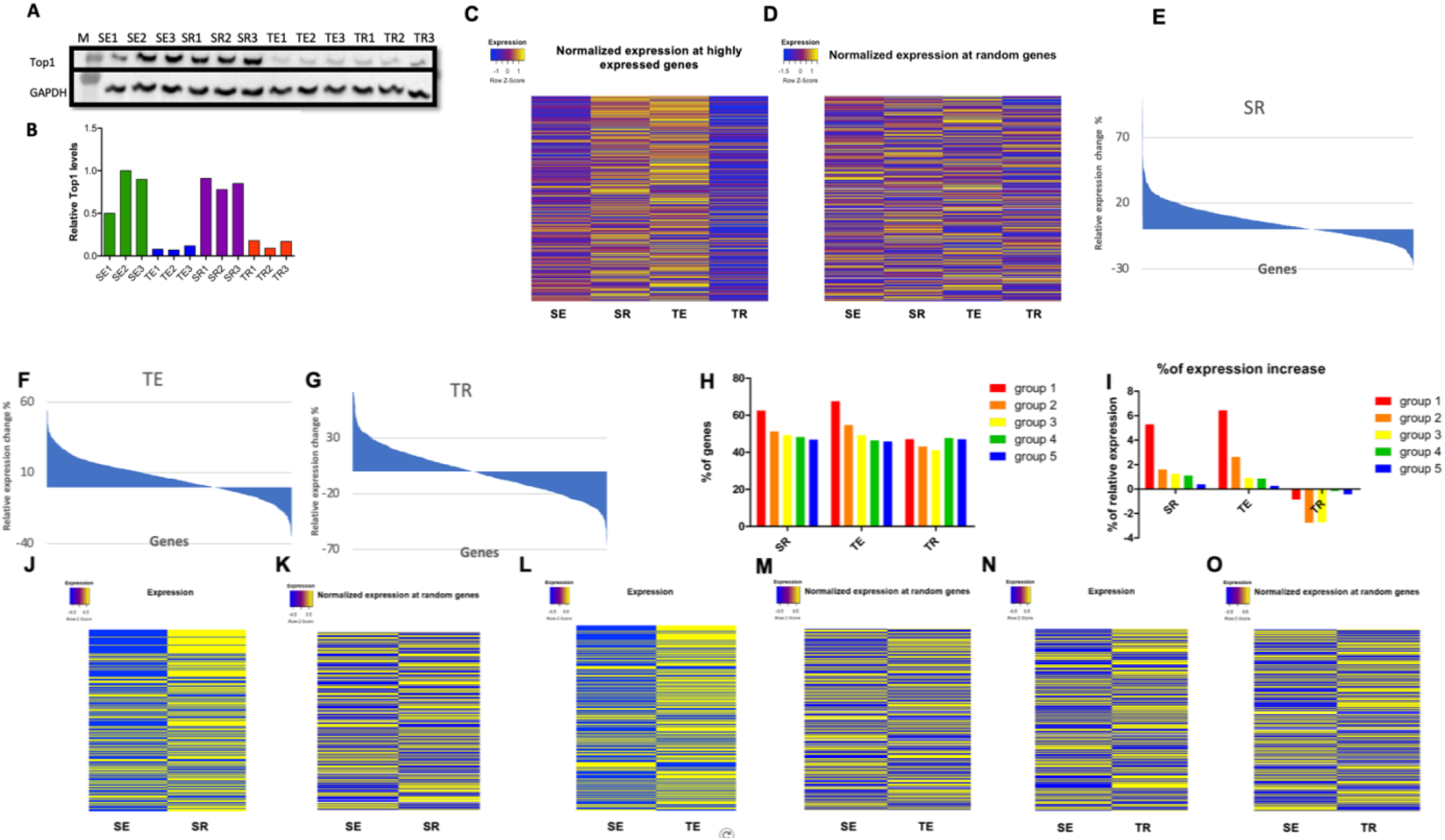
Transcriptional DSBs might be impeding transcription. **A.** Validation for TOP1 knockdown using Western Blotting. **B**. Quantification of relative TOP1 levels after TOP1 knockdown. **C-D.** color coded heatmap showing the change of normalized expression counts after TOP1 KD and/or RNase H OE at highly expressed genes (**C**) and randomly selected genes (**D**). **E-G.** Area chart showing the proportion of the genes with a decrease or increase in expression with each treatment. **H**. Percentage of genes that exhibited an increase in their normalized expression counts with RNase H1 overexpression (SR), TOP1 knockdown (TE), or both manipulations (TR), the top 5000 genes were sorted into 5 groups (1000 gene each) according to expression, the first group being the group with the top 1000 expressed genes. **I**. Percentage of normalized expression counts change with RNase H1 overexpression (SR), TOP1 knockdown (TE), or both manipulations (TR), the top 5000 genes were sorted into 5 groups (1000 gene each) according to expression, the first group being the group with the top 1000 expressed genes. **J-O**. color coded heatmap showing the change of normalized expression counts at genes with decreased breaks and randomly selected genes for reference.

**Supplementary Figure 7:**
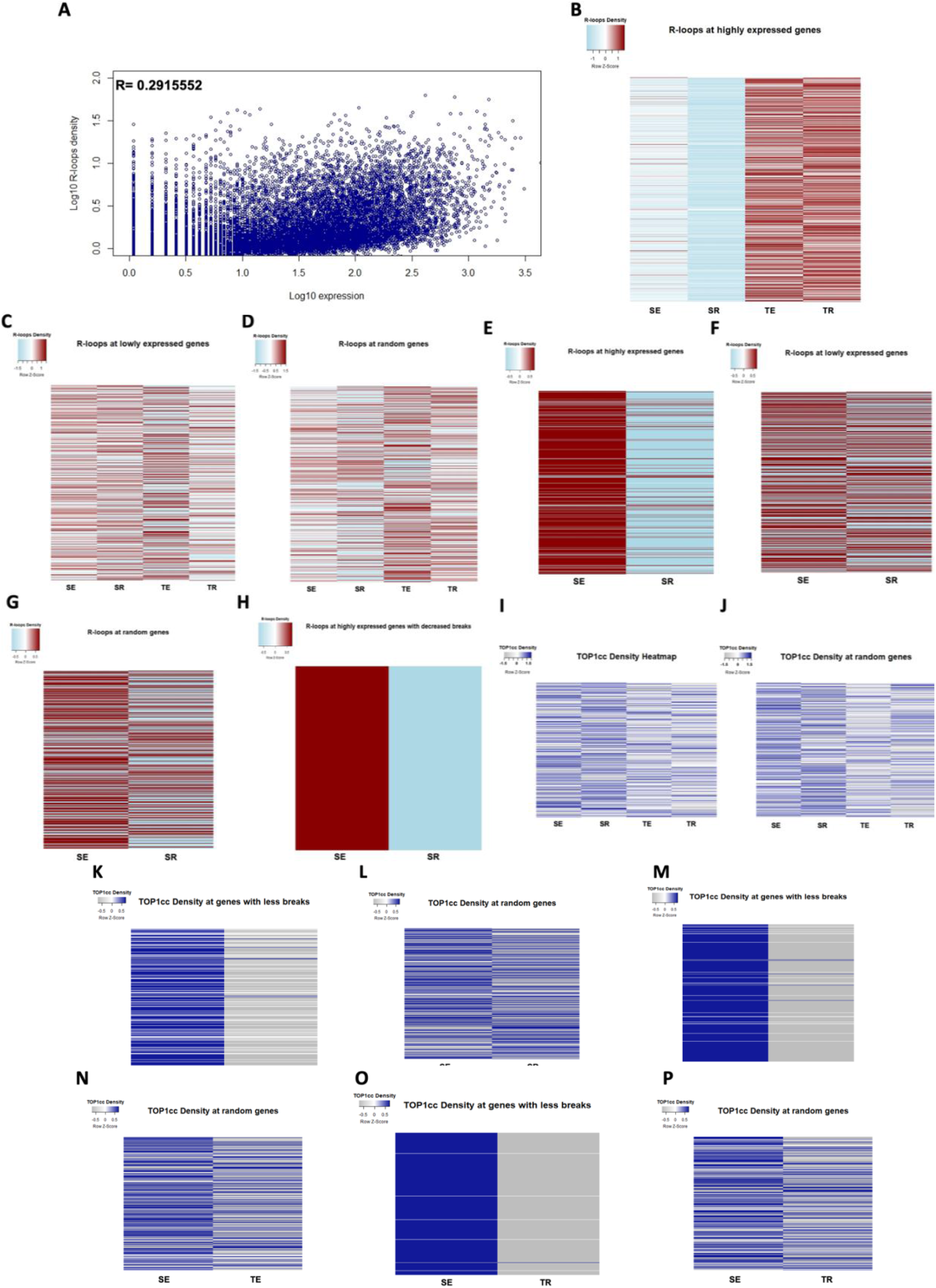
RNase H OE decreases R-loops at highly expressed genes. **A.** scatter plot showing the correlation between R-loops and transcriptional levels. **B.** color coded heatmap showing the change of DRIP signal after RNAse H OE (SR) TOP1 KD (TE) and the combined manipulation (TR) at highly expressed genes. **C.** color coded heatmap showing the change of DRIP signal after RNase H OE at lowly expressed genes. **D.** color coded heatmap showing the change of DRIP signal after RNase H OE at randomly selected genes. **E.** color coded heatmap showing the change of DRIP signal after RNase H OE at highly expressed genes. **F.** color coded heatmap showing the change of DRIP signal after RNase H OE at lowly expressed genes. **G.** color coded heatmap showing the change of DRIP signal after RNase H OE at randomly selected genes. **H.** color coded heatmap showing the change of DRIP signal after RNase H OE at highly expressed genes with decreased breaks. **I.** color coded heatmap showing TOP1cc density at lowly expressed genes. **J-P.** color coded heatmap showing TOP1cc density at the specified gene sets.

**Supplementary Figure 8:**
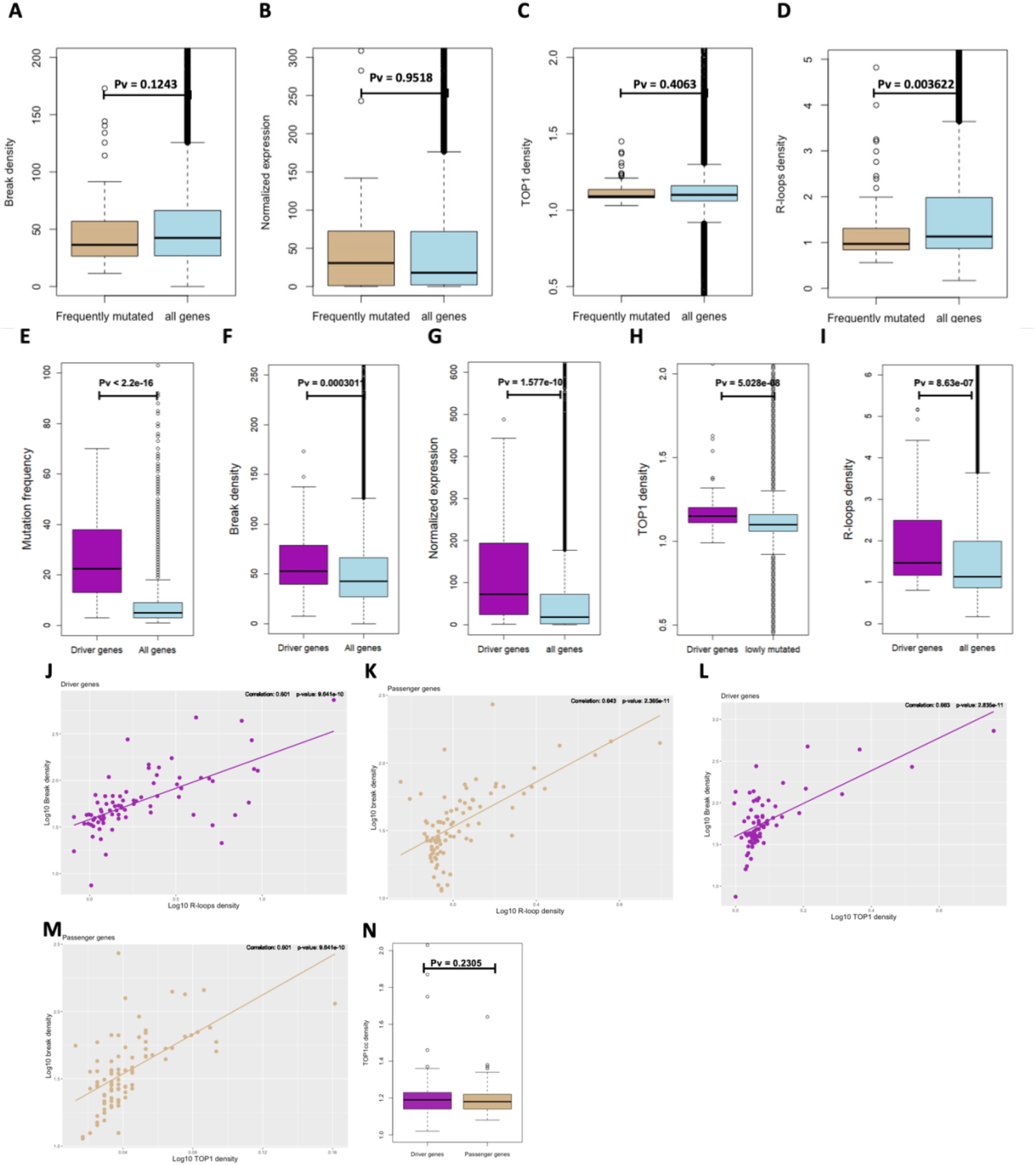
Driver genes and cancer genes are enriched with transcriptional DSBs. **A.** Boxplot of break density at frequently mutated genes and the broader gene population. **B.** Boxplot of Normalized expression at frequently mutated genes and the broader gene population. **C.** Boxplot of TOP1 density at frequently mutated genes and the broader gene population. **D.** Boxplot of R-loops density at frequently mutated genes and the broader gene population. **E.** Boxplot of mutation frequency at driver genes and the broader gene population. **F.** Boxplot of break density at driver genes and the broader gene population. **G.** Boxplot of normalized expression at driver genes and the broader gene population. **H.** Boxplot of TOP1 density at driver genes and the broader gene population. **I.** Boxplot of R-loops density at driver genes and the broader gene population. **J.** log-scaled scatter plot demonstrating the correlation between break density and R-loops density at driver genes. **K.** log-scaled scatter plot demonstrating the correlation between break density and R-loops density at passenger genes. **L.** log-scaled scatter plot demonstrating the correlation between break density and TOP1 density at driver genes. **M.** log-scaled scatter plot demonstrating the correlation between break density and TOP1 density at passenger genes. **N.** Boxplot of TOP1cc density at driver genes and passenger genes.

**Supplementary Figure 9:**
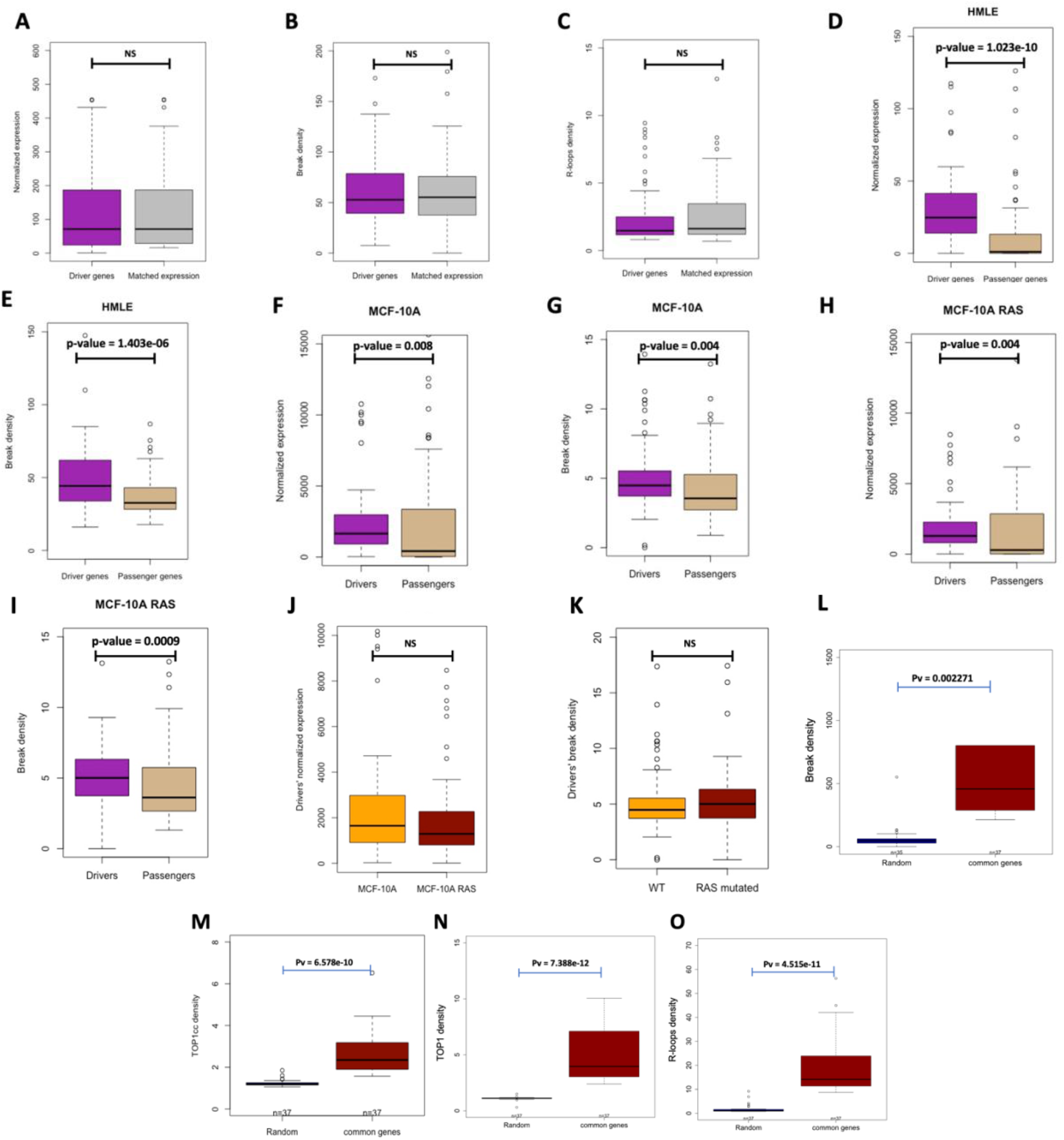
physiological DSBs as an early event of carcinogenesis. **A.** Boxplot of normalized expression at driver genes and genes matched for expression. **B.** Boxplot of break density at driver genes and genes matched for expression. **C.** Boxplot of R-loops density at driver genes and genes matched for expression. **D.** Boxplot of normalized expression of driver genes and passenger genes in HMLE cells. **E.** Boxplot of break density of driver genes and passenger genes in HMLE cells. **D.** Boxplot of normalized expression of driver genes and passenger genes in MCF-10A cells. **E.** Boxplot of break density of driver genes and passenger genes in MCF-10A cells. **F.** Boxplot of normalized expression of driver genes and passenger genes in MCF-10A cells. **G.** Boxplot of break density of driver genes and passenger genes in MCF-10A cells. **H.** Boxplot of normalized expression of driver genes and passenger genes in RAS mutated MCF-10A cells. **I.** Boxplot of break density of driver genes and passenger genes in RAS mutated MCF-10A cells. **J.** Driver genes normalized expression in MCF-10A and RAS mutated MCF-10A cells. **K.** Driver genes break density in MCF-10A and RAS mutated MCF-10A cells. **L-O.** Boxplot of break density (**L**), TOP1cc density **(M)**, TOP1 density (**N**), and R-loops density (**O**) at intersected genes of Figure 7Q and randomly selected genes. **S.** Boxplot comparing driver genes to non-driver genes matched for expression in expression, break density, and R-loops respectively.

## Notes

### Competing Interest Statement

The authors have declared no competing interest.

### Summary of Updates

Detailed analysis of breakomes in MCF7, MCF10A and HMLE cells establishing a link between transcriptional DSBs and early molecular changes driving cancer development. Notably, our study highlights the distinct etiology and molecular characteristics of driver mutations compared to passenger mutations, shedding light on the potential for targeted therapeutic strategies.

